# The novel RNA polymerase I transcription inhibitor PMR-116 exploits a critical therapeutic vulnerability in a broad-spectrum of high *MYC* malignancies

**DOI:** 10.1101/2025.04.19.649466

**Authors:** Rita Ferreira, Katherine M. Hannan, Konstantin Panov, Thejaani Udumanne, Amee J. George, Adria Closa, Zaka Yuen, Eduardo Eyras, Perlita Poh, Kylee Maclachlan, Mitchell Dwyer, Vanessa Barn, Jinshu He, Eric Kusnadi, Richard Rebello, Alisee Huglo, Shelley Hedwards, Mitchell Lawrence, Gail Risbridger, Renea Taylor, Ashlee Clark, Geoffrey Farrell, Sharon Pok, Narci Teoh, Madushani Amarasini, Philip L. Norcott, Martin G. Banwell, Yulia Manenkova, Mustapha Haddach, Denis Drygin, Luc Furic, Ross D. Hannan, Nadine Hein

## Abstract

Ribosome biogenesis (RiBi) is a key determinant of cell growth and proliferation and is highly elevated in cancer due to the activation by oncogenes such as *MYC*. First-generation RiBi inhibitor CX-5461, while demonstrating clinical potential for cancer treatment, also induces DNA damage through off-target inhibition of TOP2⍺ and potentially other mechanisms, bringing into question RiBi as a target for cancer therapy. In this study, we test second-generation RiBi inhibitor, PMR-116. PMR-116 exhibits improved drug-like properties compared to first-generation RiBi inhibitors and has robust anti-tumour activity in the absence of global DNA damage signalling in a broad range of pre-clinical models of haematologic and solid cancers, particularly in malignancies where *MYC* is either the driver of disease or is elevated. Thus, our work demonstrates that RiBi is a genuine target for cancer therapy and highlights the potential to exploit a critical therapeutic vulnerability in high-*MYC* human cancers with dismal therapeutic outcomes.

**Statement of significance:** Despite the development of new cancer therapies, most advanced malignancies remain incurable. We demonstrate that PMR-116, a second-generation RiBi inhibitor, has robust therapeutic efficacy in preclinical models of cancer, offering great promise to treat a broad spectrum of human solid and haematologic malignancies, especially where *MYC* is a driver.

## Introduction

Ribosome biogenesis (RiBi) is a highly conserved process that occurs in the nucleolus and controls cellular growth and proliferation [1]. Transcription of ribosomal RNA (rRNA) genes (rDNA) by RNA Polymerase I (Pol I) is the first critical step in RiBi and is tightly regulated by tumour suppressors such as RB, p53 and PTEN, and oncoproteins such as MYC which together regulate either directly and indirectly the Pol I machinery to tightly control proliferation in non-cancerous cells [2–4]. During malignancy, loss of function of tumour suppressors and upregulation of oncogenes lead to a significant elevation in Pol I transcription, in part due to the robust increase in expression of Pol I subunits, associated transcription factors and the activation of previously silenced rRNA genes [3–8]. Moreover, the potent oncogenic activity of MYC, which is amplified in 70% of human cancers, can at least in part be explained by its ability to drive RiBi [9, 10].

Together, these data provide a strong rational for the targeting of key regulatory steps in RiBi to treat malignancies, especially those driven by MYC [5, 9, 11]. The development of the first-generation small molecule inhibitor of RNA Polymerase I transcription, CX-5461 [12, 13] provides further support for this; demonstrating strong preclinical activity in multiple models of cancer [13–18]. Additionally, Phase I clinical trials with CX-5461 in patients with haematological and solid tumours also exhibit promising initial clinical activity [19, 20]. Concurrent with these studies, CX-5461 was identified to induce a DNA damage response (DDR) [17, 21, 22], which is not predicted from the pathways known to be activated downstream of Pol I transcription inhibition, most notably the p53 dependent and independent nucleolar surveillance pathways (NSP) [5], and is not observed for other Pol I inhibitors such as BMH-21 [23]. A number of studies propose additional mechanisms of action (MOAs) for CX-5461 to those originally reported [24–29], including TOP2α poisoning [27, 29, 30]. Concerningly, a recent study by Koh *et al*. provided evidence that CX-5461 can cause rapid, extensive, and non-selective mutagenesis in cultured cell lines to magnitudes that exceed known environmental carcinogens and clinically used chemotherapeutic drugs [31]. Through genetic approaches, we have recently resolved this conundrum, proving that, in fact, CX-5461 dually, but independently target RNA Polymerase I and TOP2α activity [30].

Thus, it remains unclear whether directly targeting Pol transcription is a viable and efficacious approach to treating cancer in the absence of DNA damage and its associated problems, such as cardiomyopathy and the induction of secondary cancers. To clarify this matter, we developed a series of more selective second-generation Pol I inhibitors with our lead compound being PMR-116. We demonstrate that PMR-116 exhibits improved drug-like properties and pharmacokinetics compared to CX-5461, and has robust anti-cancer activity in a range of syngeneic and xenograft preclinical models of haematologic and solid tumours with high potency in those driven by MYC. Critically, the therapeutic response to PMR-116 occurs in the absence of DNA damage signalling, thus establishing inhibition of Pol I transcription as a *bone fide* and promising therapeutic strategy for cancer therapy, especially where MYC is a primary driver of disease.

## Results

### PMR-116 selectively inhibits rRNA synthesis by stalling RNA Polymerase I at the rDNA promoter regions

The second-generation Pol I transcription inhibitor PMR-116 was developed by Pimera Therapeutics using a phenotypic screening approach (qRT-PCR based assay) as described previously [12]. The core molecular structure of PMR-116, unlike CX-5461, contains a non-quinolone moiety, as presented in **Figure 1A**. The potential of PMR-116 to inhibit Pol I transcription was confirmed in cell-free transcription assays using purified Pol I complex components. PMR-116 inhibited Pol I specific promoter-driven transcription with higher sensitivity compared to randomly initiated (non-specific) transcription with a mean transcription inhibitory concentration (TIC) 50 of 19 μM and 528 μM, respectively (**Figure 1B**). The differential selectivity of PMR-116 to suppress Pol I over Pol II transcription was determined *in vitro* in primary human cells and cancer cell lines. This was assessed by measuring the rapidly post-transcriptionally processed 5’ETS region of the Pol I transcribed 47S pre-rRNA [32] and the Pol II transcribed cFOS and MYC mRNAs (**Figure 1C & SFigure 1A**). PMR-116 inhibited rRNA synthesis with mean TIC50s between 0.08-0.59 μM, while the TIC50s for Pol II-dependent transcription were at least two orders of magnitude higher (mean Pol II TIC50s >3 or >25 μM), thus confirming PMR-116 as selective Pol I transcription inhibitor. It is important to recognise that fully replicating cellular conditions in a reconstituted transcription assay is not feasible. Therefore, significant differences between cell-based and cell-free assays are expected.

**Figure 1.**
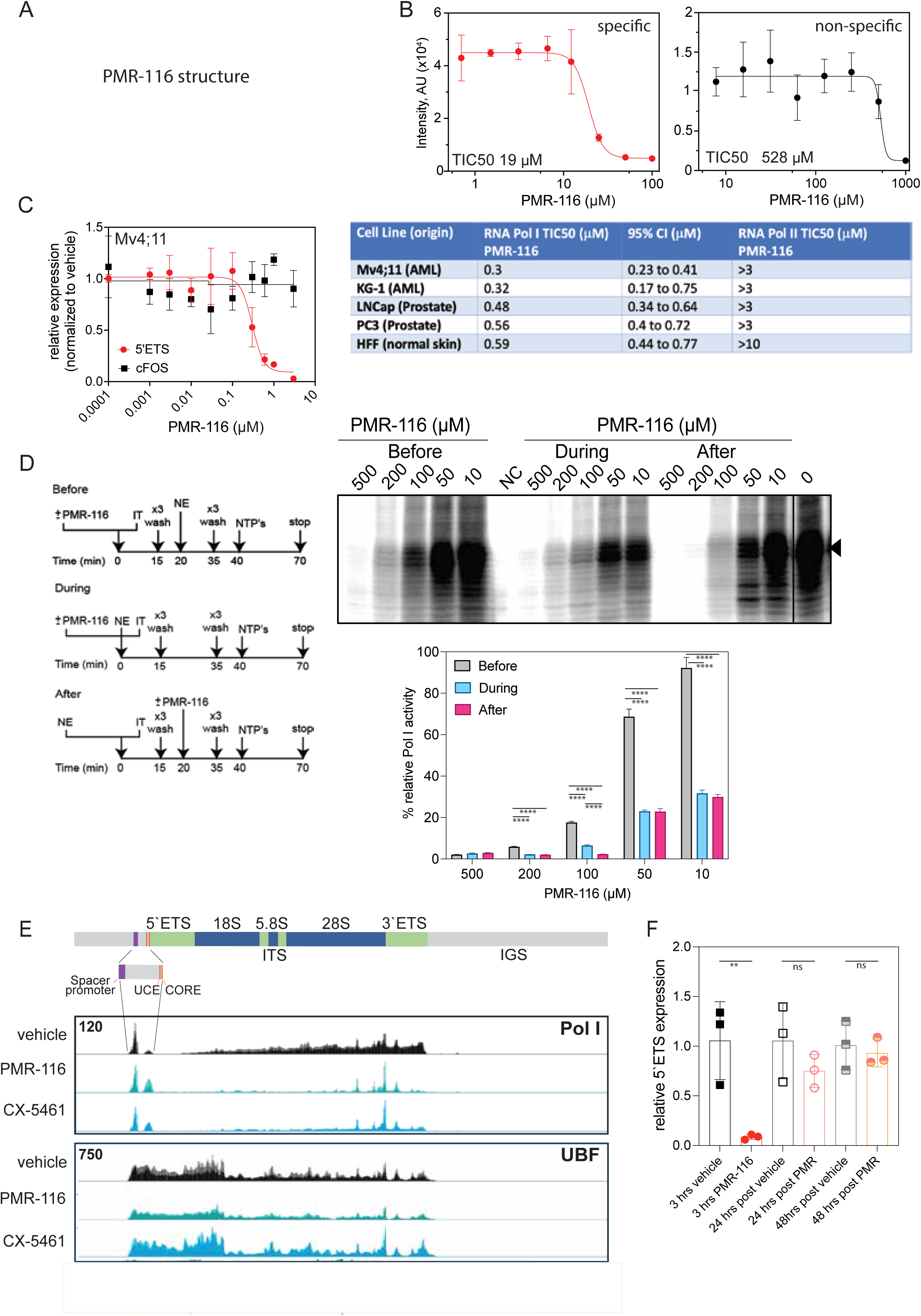
Second generation Pol I inhibitor PMR-116 selectively inhibits RNA polymerase I transcription. **A** Chemical structure of PMR-116 **B** In vitro PMR-116 inhibits promotor-driven (specific) Pol I transcription with a TIC50 of 19 µM vs randomly initiated (non-specific) Pol I transcription with a TIC50 of 528 µM. The dose-response curve and TIC50 was plotted and calculate in GraphPad Prism 10 using non-linear regression ([inhibitor] versus response – variable slope (four parameters)). The data represent the mean -/+ SD of three independent biological experiments (n=3). **C** Relative expression of 5’ETS (Pol I) and cFOS (Pol II) assessed by qPCR in Mv4;11 cells after 3 hrs of treatment with increasing concentrations of PMR-116 (left) (n=3); average TIC50 concentration for Pol I and Pol II transcription inhibition in various human AML, prostate cancer cells lines and non-transformed fibroblasts. TIC50 were calculated as in **A** from at least three independent biological replicates (n=3). **D** Schematic overview of *in vitro* order of addition experiments (left). Representative image of order of addition assays. PMR-116 reduces in vitro transcription in a dose dependent manner when added during or after PIC formation. Quantification of relative Pol I activity in order of addition experiments presented as the mean percentage ± SD (n=3) (lower panel); ordinary one-way ANOVA, *P* **** <0.0001; NE – nuclear extract; IT-immobilized template. **E** schematic presentation of a single rRNA gene (upper panel). IGV plots visualise Pol I (RPA194) and UBF enrichment at rRNA gene after treatment with either vehicle, 1 µM PMR-116 or 1 µM CX-5461 in Mv4;11 cells 30min post treatment, overlay of n=3 independent biological replicates.

The mechanism by which PMR-116 interferes with Pol I activity was determined utilising cell-free transcription and Chromatin Immunoprecipitation (ChIP) assays. The Pol I transcription cycle begins with the assembly of a productive preinitiation complex (PIC) at the rDNA promoter, followed by promoter-escape and transcription elongation stages. PMR-116 efficiently repressed Pol I transcription in reconstituted transcription reactions supplemented with nuclear extracts or with partially purified Pol I transcription components (**SFigure 1B**) [33] suggesting that PMR-116 directly inhibits Pol I transcription either at the initial stages of transcription (such as PIC assembly or promoter-escape), or affects the catalytic activity of Pol I. The latter is unlikely as PMR-116 does not inhibit non-specific Pol I transcription, excluding Pol I elongation and thus catalytic activity as the main PMR-116 target (**Figure 1B**). To test whether PMR-116 inhibits Pol I transcription PIC formation (**SFigure 1C**) or promoter-escape, order of addition transcription reactions were performed using an immobilized template (IT) containing the Pol I promoter nuclear extracts (NEs) (**Figure 1D**) [11]. Transcription efficiency was analysed when PMR-116 was added before, during or after the addition of NEs. Pol I transcription was most efficiently inhibited when the drug was added during or after PIC formation, suggesting that it likely does not bind to the promoter in the absence of the PIC but inhibits the formation of productive PIC and/or interferes with post PIC formation events such as promoter clearance (**Figure 1D**). Consistent with PMR-116 affecting promoter clearance, ChIP-sequencing (seq) for RPA194, the largest subunit of Pol I, in human Mv4;11 AML cells treated with PMR-116 (3x TIC50) demonstrated an almost complete depletion of Pol I loading within the transcribed region of the rDNA coding region while Pol I enrichment at the upstream control element (UCE) regions of the promoter remained unaffected (**Figure 1E).** These changes were similar to those observed in response to CX-5461 treatment. Consistent with decreased Pol I occupancy in the transcribed region following PMR-116 treatment, the enrichment of the Pol I specific polymerase-associated factor (PAF) 53 was also reduced in a pattern similar to that of Pol I (**SFigure 1D**). The Pol-I specific cytoarchitectural transcription factor Upstream Binding Factor (UBF), demonstrated reduced loading in the UCE and transcribed regions in response to 1 μM PMR-116 but remained unaffected by CX-5461 treatment (**Figure 1E**). In marked contrast to CX-5461, which permanently blocks Pol I initiation and thus Pol I transcription [22], PMR-116 mediated Pol I inhibition is reversible, i.e. Pol I transcription is restored when PMR-116 is washed out from cells (**Figure 1F**).

We also tested the ability of PMR-116 to affect the formation of G quadruplex (G4) structures which have the potential to inhibit rRNA synthesis [34]. In contrast to TMPyP4, a known G4 stabilizer [35], PMR-116 did not significantly increase G4 structures, as detected by immunofluorescence in Mv4;11 and KG-1 cells (**SFigure 1E**). Interestingly, PMR-116 treatment at 3xTIC50 concentration led to degradation of the large Pol I subunit (RPA194) in KG-1, similar to the DNA intercalator BMH-21 [36], but not in Mv4;11 cells (**SFigure 1F**). Therefore, we conclude that at selective on-target concentrations, PMR-116 inhibits Pol I transcription by enhancer/promoter stalling of Pol I thus preventing promoter escape.

### PMR-116 has improved drug-like properties compared to CX-5461 and possesses broad anti-cancer activity *in vitro*

The pre-selection and development criteria for an improved 2^nd^ generation Pol I transcription inhibitor over 1^st^ generation Pol I inhibitor CX-5461 were: 1) improved pharmacokinetic properties (higher bioavailability); 2) better solubility 3) potential to penetrate the blood-brain barrier and 4) limited induction of DNA damage signalling. Consistent with these criteria, PMR-116 outperforms CX-5461 with respect to their drug-like properties and pharmacokinetics (**Figure 2A**). PMR-116 has a lower molecular weight (388.5 g/mol vs 513 g/mol), higher solubility (360 mg/ml vs 25 mg/ml), and increased bioavailability (82% vs 24% in mice) [13]. The potential to cross the blood-brain barrier (BBB) was tested *in vivo* with a single dose of 10 mg/kg PMR-116 administered by oral gavage, achieving around 18-fold higher drug concentration in the brain than plasma (**Figure 2B**). PMR-116 exhibited a mean C_max_ value of 23094.37 ng/g in the brain vs 1274.66 ng/ml in the plasma, displayed a moderate half-life of 9.33 hrs in the brain and 16.82 hrs in the plasma and demonstrated favourable systemic exposure with an area under the curve (AUC) _0-infinity_ of 281577.5 ng/g*hr in the brain and 20390 ng/ml*hr in the plasma. The high brain concentration achieved is likely due to the PMR-116’s LogPe (the PAMPA effective permeability coefficient) value of -5.4 ± 0.1, which is comparable to the LogPe value for an approved CNS-targeting therapeutic Propranolol (-5.3 ± 0.1) as measured by Parallel Artificial Membrane Permeation Assay (PAMPA) modified to mimic the BBB (**Figure 2B)** [37].

**Figure 2.**
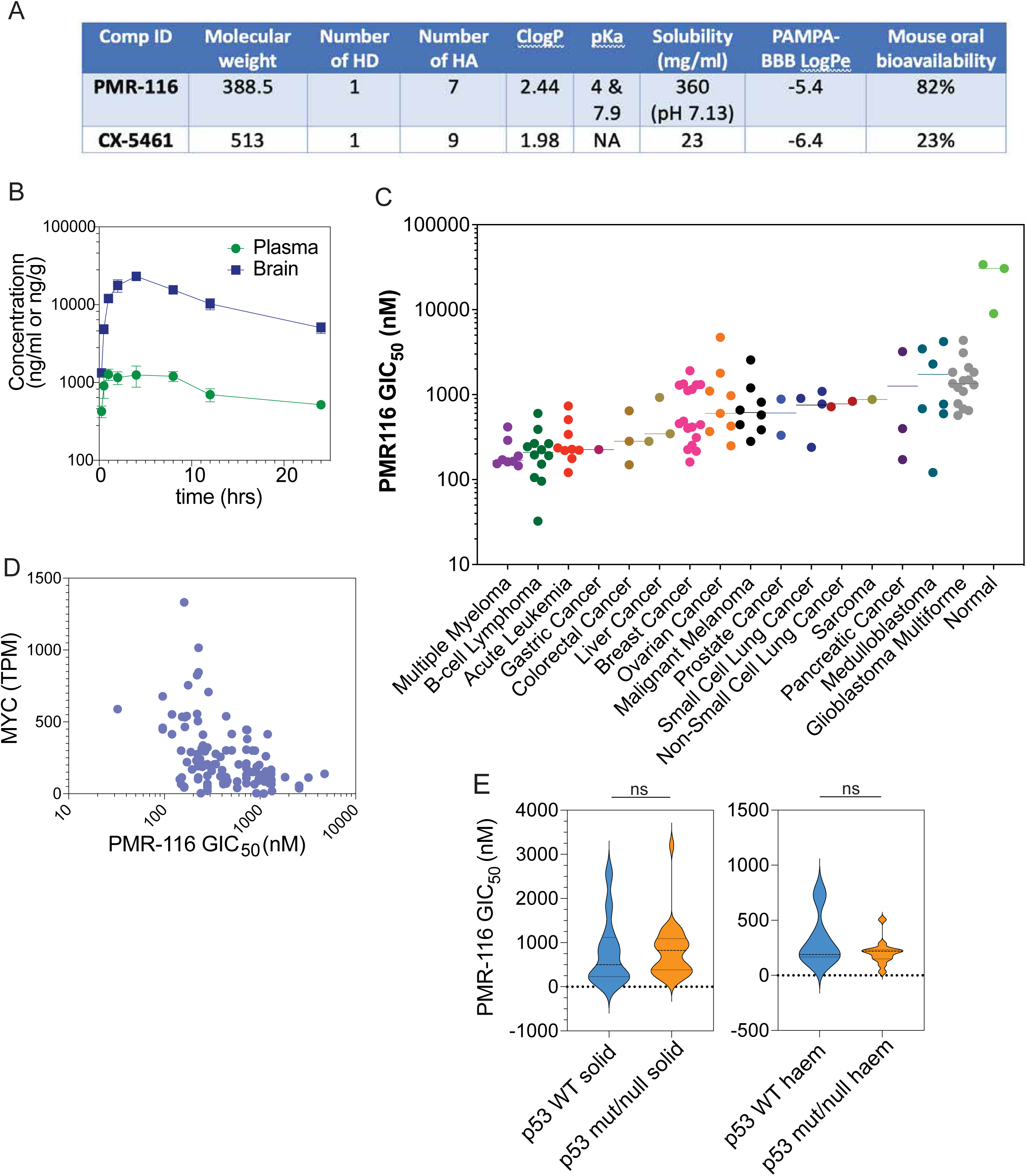
PMR-116 displays improved drug-like properties and has anti-cancer activity in human – derived cancer cell lines. **A** Comparison of chemical and pharmacokinetic properties of PMR-116 and CX-5461. **B** PMR-116 concentration in the plasma and brain in mice treated with a single dose of 100 mg/kg PMR-116 via oral gavage. **C** The average GIC50 was assessed after 96 hrs of treatment with PMR-116 in a panel of 107 human-derived haematological and solid tumour cancer cell lines and normal cells. The average GIC_50_ was determined using non-linear regression, sigmoidal dose-response in GraphPad Prism v10 (n=1-20 per cell line). **D** Relationship between MYC expression level plotted as transcripts per million (TPM) and PMR-116 GIC_50,_ non-parametric Spearman correlation r = -0.4815, (95% CI -0.6523 to -0.2643, *P* <0.0001, two-tailed) **E** The violin plots display the average GIC_50_ in p53WT and mutant/null haematological (right) and solid (left) tumours. The p53 status does not determine sensitivity to PMR-116 induced growth inhibition, two-tailed unpaired t-test, ns = non-significant.

While PMR-116 has superior drug-like properties over first-generation CX-5461, PMR-116 also displayed a broad spectrum of anti-proliferation activity (150-5000 nM a > 30 fold range) across a panel of human and murine cancer cell lines with an median growth IC50 (GIC50) of approximately ∼470 nM, while normal cells were significantly more resistant to PMR-116 with an median GIC50 ∼60 fold higher (∼30 μM). Blood-derived tumours (myeloma, B-cell lymphoma) were the most sensitive, followed by liver, colorectal, ovarian, prostate and breast cancer (**Figure 2C**). Intriguingly, cell sensitivity was found to inversely correlate with MYC expression (CI -0.6523 to -0.2643, -0.4815, p < 0.0001) (**Figure 2D and SFigure 2A**) but was largely independent of the p53 status in both haematological and solid cancers (**Figure 2E)**. The difference in sensitivity of non-cancerous vs malignant cell lines to PMR-116’s growth inhibitory effect was not due to differences in the Pol I TIC50s as both the breast adenocarcinoma cell line MDA-MBA-231 and CCD-1096Sk, a fibroblast cell line established from normal breast skin, displayed similar TIC50s of 118 and 79 nM **(SFigure 2B)**. However, the GIC50s for these cell lines differed greatly, 141 and 30,500 nM, respectively (**SFigure 2B**). These data indicate that PMR-116 has the potential to act as a broad-spectrum anti-cancer drug with its sensitivity linked to high expression of MYC, but independently of p53 status.

### Inhibition of Pol I transcription induces global transcriptional changes consistent with activation of nucleolar surveillance pathways

To identify signalling pathways altered by PMR-116 treatment that mediate its growth inhibitory phenotype and to distinguish differentially regulated pathway compared to CX-5461 transcriptome profiling was performed in Mv4;11 and KG-1 treated with both inhibitors. PMR-116 treatment induced robust changes in the global transcriptome in Mv4;11 while p53 null KG-1 displayed considerably less changes at both timepoints (3hrs: 4222 vs 1811 differentially expressed genes (DEG); 12hrs: 7763 vs 1793 DEG) (**Figure 3A&E**). Of note, the same number of genes were upregulated or downregulated at both timepoints, consistent with **Fig 1** showing that PMR-116 does not inhibit Pol II transcription at TIC90 concentrations, as one might expect a majority of downregulated genes if this was the case. Interestingly PMR-116 treatment induced more widespread transcriptional changes compared to CX-5461 treatment (**Figure 3, SFigure 3**) with <50% of DE genes overlapping between both drugs in Mv4;11 cells and less <20% overlap observed in KG-1 cells at both timepoints (<20%) (**Figure 3B&F**).

**Figure 3.**
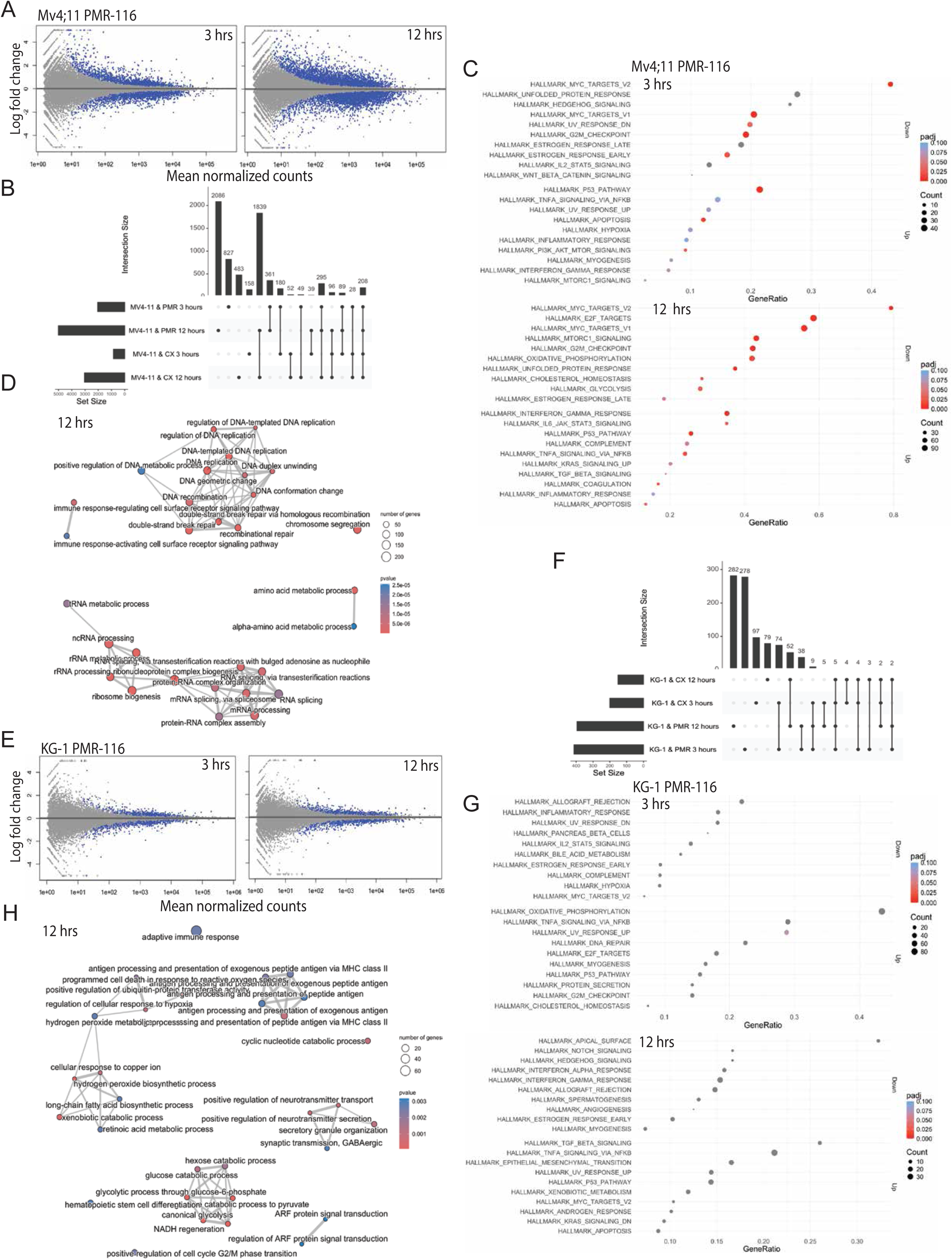
Pol I transcription inhibition induces global transcriptional changes. **A** MA plots show normalized read counts between vehicle and PMR-116 treated Mv4;11 cells, plots visualize significant differentially expressed (DE) genes (blue dots) at 3 and 12 hrs post PMR-116 treatment (3 x TIC90) compared to vehicle (upper panel). The plots show significant DE genes identified in all three independent biological replicates **B** Upset plot visualises the intersection of DE genes between all treatment groups and timepoints in Mv4;11 cells. Set size indicates the number of DE genes in each group and the intersection the number of DE genes in different groups that overlap. **C** Hallmark pathway analysis in Mv4;11 at both timepoints in response to PMR-116 treatment. **D** Enrichment Map analysis organised the top 30 deregulated pathways based on their overlapping gene sets into functional module (Mv4;11; 12 hrs). **E** MA plots showing significant DE genes (blue dots) in KG-1cells after 3 and 12 hrs of PMR-116 (3 x TIC90) treatment (n=3 independent biological replicates per timepoint). **F** Upset plots displays the number of overlapping DE genes between the different treatments and timepoints in KG-1 cells. **G** Hallmark pathway analysis in KG-1 cell at 3 and 12 hrs of treatment. **H** Enrichment Map analysis displays functional hubs differentially regulated after 12 hrs of PMR-116 treatment in KG-1 cells.

Hallmark pathway analysis after 3hrs of PMR-116 treatment in MV4;11 cells exhibited significant upregulation of p53 target genes and apoptosis-related genes, alongside downregulation of MYC target genes, G2M checkpoint genes and genes involved in UV response (**Figure 3C**). After 12 hrs, additional significant Hallmark pathway changes were observed, including upregulation of interferon-gamma response pathways, TNF⍺ signalling via NF*κ*B, and downregulation of MYC and E2F targets, mTORC1 signalling, and genes involved in oxidative phosphorylation, and glycolysis. Critically, modulation of p53, MYC, E2F, mTORC, G2M checkpoint genes, and NF*κ*B signalling are well-described targets of the canonical and non-canonical NSP following inhibition of Pol I transcription and RiBi; hence, they are consistent and likely major mediators of the therapeutic response to PMR-116 treatment (**Figure 3B**) [38]. The downregulation of MYC target genes is consistently observed in Mv4;11 cells treated with either PMR-116 or CX-5461, further highlighting the potential to target aberrant MYC target gene expression signatures with Pol I inhibitors (**Figure 3C, SFigure 3B**). Enrichment map analysis after 12 hrs treatment with PMR-116 revealed two deregulated gene networks, RiBi and mRNA-associated processes as well as DNA replication and DNA recombination/repair, whereas CX-5461 treatment mainly affected gene networks linked to DNA repair, chromosome segregation and metabolic processes (**Figure 3C, SFigure 3C**). In line with the modest overall transcriptional changes, Hallmark pathways and networks enriched in p53null KG-1 in response to PMR-116 and CX-5461 exposure, while not statistically significant, were consistent with non-canonical NSP response pathways, including MYC pathways (**Figure 3E-G, SFigure 3D-F**). In conclusion, PMR-116 induces transcriptional programs consistent with the p53-dependent canonical and p53-independent non-canonical NSP.

### PMR-116 increases p53 protein abundance and delays cell cycle progression but does not activate DNA damage signalling

Previous studies have demonstrated that acute inhibition of Pol I transcription activates the NSP resulting in the accumulation of the tumour suppressor protein p53 and subsequent induction of downstream responses such as apoptosis, senescence and/or cell cycle arrest [5, 16]. Consistent with this and the transcriptomic analysis, PMR-116 treatment of Mv4;11 (p53WT) at 3x Pol I TIC50 concentrations induced modest apoptotic cell death at 3xTIC50 concentrations (**Figure 4A**; late apoptosis: vehicle 0.65%±0.05% vs 3xTIC50 7.65% ±3.04%; early apoptosis: vehicle 1.36% ±0.06% vs 3xTIC50 6.72%±2.27%), and a significant decrease in S phase cell populations (**Figure 4B**; vehicle 30.78%±1.08% vs TIC50 3.45%± 2.89% vs 3xTIC50 2.48% ±0.98%). In the p53null AML cell line KG-1 no significant apoptotic cell death was detected but a decrease of cells in S-phase (vehicle 35.53% ±8.35% vs TIC50 23.92%±1.73% vs 3xTIC50 24.89%±3.6) and a significant increase in the G2 cell population (**Figure 4A & B;** vehicle 8.57%±2.46 vs TIC50 12.09% ±1.6% vs 3xTIC50 24.91% ±2.04%) was observed after 72 hrs of treatment. The human prostate cancer cell line LNCaP underwent apoptotic cell death in response to Pol I transcription inhibition in contrast to PC3. Both displayed aberrant cell cycle progression with LNCaP cells exiting S-phase and PC3 cells accumulating in G2/M phase of the cell cycle (vehicle 9.47% ±0.17% vs TIC50 18.53%±3.08% vs 3xTIC50 34.86%±2.6) (**Figure 4A & B**). Concomitant with the induction of nucleolar stress, PMR-116 treatment (3xTIC50) disrupted nucleolar morphology and induced nucleolar dispersion and cap formation in Mv4;11 and KG-1 cells (**Figure 4C & SFigure 4A**). Additionally, PMR-116 increased p53 protein abundance in p53 WT cancer cell lines, Mv4;11, LNCaP and non-cancerous fibroblasts (**Figure 4D & E**). Consistent with previous studies CX-5461 treatment (3x Pol I TIC50) led to phosphorylation of p53 at Serine (Ser) 15, a modification induced following DNA damage that modulates p53 transactivation potential and activation of the p53 target gene p21 in p53WT cell lines Mv4;11and LNCaP. In contrast, both p53-Ser15 and p21 were not induced in Mv4;11 treated with 3x TIC50 concentration of PMR-116 (**Figure 4D**).

**Figure 4.**
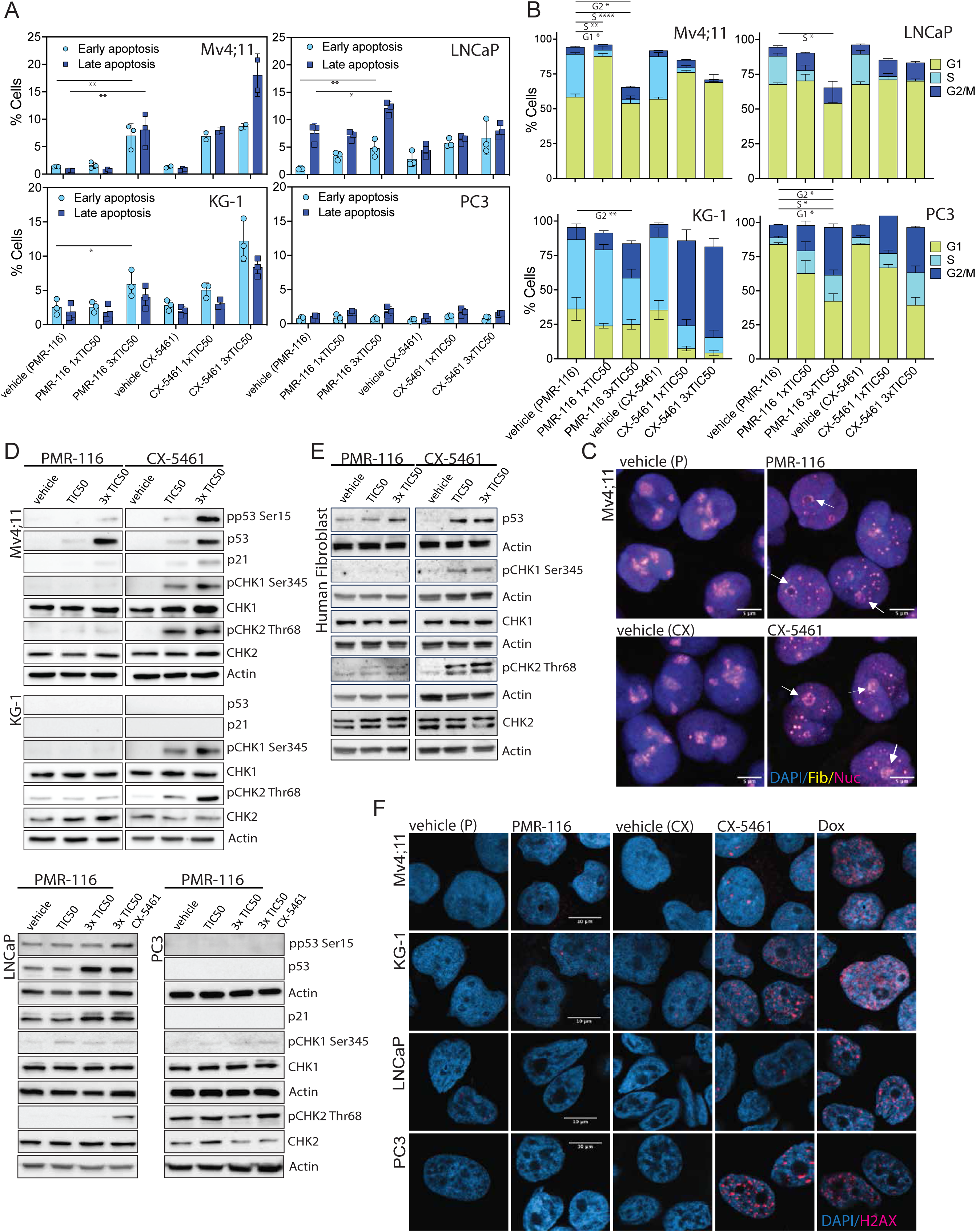
Phenotypic response and pathway activation in response to PMR-116 in AML and prostate cancer cell lines. **A** Apoptotic cell death was analysed by AnnexinV/7AAD staining after 48 hrs of treatment with either vehicle, TIC50 or 3xTIC90 concentration of either PMR-116 or CX-5461 in Mv4;11, KG-1, LNCap and PC3 cells. Cells were categorized as late or early apoptotic by being AnnexinV and 7AAD double positive or AnnexinV positive respectively. Bar graphs represent the mean ± SD of n=2-3; *P* ** <0.009, *P* * <0.05; ordinary one-way ANOVA, Dunnett’s multiple comparison. **B** Cell cycle distribution was assessed by BrdU incorporation and PI staining 72 hrs post treatment with either vehicle, TIC50 or 3xTIC90 concentration of either PMR-116 or CX-5461 in Mv4;11, KG-1, LNCaP and PC3 cells. Cell population in G0/G1, S and G2/M were distinguished based on BrdU incorporation and DNA content. The graphs represent the mean ± SD of n=3; two-way ANOVA Tukey’s multiple comparisons *P* **** <0.0001, *P* ** <0.007, *P* * <0.05. **C** Fibrillarin (yellow) and Nucleolin (pink) immunofluorescent staining in Mv4;11 cells treated for 3 hrs with either vehicle (citric acid or NaH_2_PO_4_), PMR-116 (3xTIC50) or CX-5461 was conducted to analyse changes in nucleolar morphology. The nucleus was stained using DAPI (blue). Images were taken at 64x magnification, scale bar represent 5µm. **D** and **E** Mv4;11, KG-1, PC3, LNCaP and primary human fibroblasts were treated with PMR-116 and CX-5461 at TIC50 and 3x TIC50 concentration for 3 hrs. Changes in p53, p21, CHK1/2 protein abundance and phosphorylation of p53-Ser15, CHK1-Ser345 and CHK2 Thr68 by immunoblotting. Actin was detected as loading control. **F** In contrast to Doxorubicin and CX-5461 in KG-1 and PC3, PMR-116 does not induce global *γ*H2AX (pink) foci in all cell lines analysed by immunofluorescence. DAPI (blue) marks the nucleus. Representative IF and immunoblot images are shown of at least n=3 biological replicates (**D-F**).

Most critical for therapeutics is the minimization of any off-target effects that may amplify unwanted side effects. CX-5461 has been shown to induce DNA damage signalling most likely via inhibition of Top2⍺. In line with our previous findings at 1x and 3x TIC50 concentrations, CX-5641 treatment led to robust phosphorylation of checkpoint kinases 1 and 2 (CHK1 and CHK2) at Ser 345 and Thr64, which lie downstream of the DNA damage sensing ATM and ATR kinases, respectively, and independent of functional p53 [16, 17]. In sharp contrast and despite the enrichment analysis revealing DNA repair linked pathways, PMR-116 did not induce phosphorylation of CHK1 and 2 above background level in any of the cell lines tested, suggesting that PMR-116 therapeutic action is independent of DNA damage (**Figure 4D**). Similarly, PMR-116 treatment at both concentrations led to only minimal, if any, phosphorylation of CHK1 and CHK2 in normal fibroblasts (**Figure 4E**). In line with absence of CHK activation is the lack of above background *γ*H2AX foci formation in response to PMR-116, while CX-5461 in KG-1 and PC3 cells and Doxorubicin, a potent DNA damaging agent, caused global *γ*H2AX foci formation after 24 hrs of treatment in all cell lines tested (**Figure 4F & SFigure 4B)**. In summary, PMR-116 treatment mediates its anti-cancer activity by altering cell cycle progression and induction of the canonical and non-canonical NSP in the absence of global DNA damage.

### PMR-116 therapy treats aggressive haematological malignancies

To explore whether PMR-116 *in vitro* anti-cancer activity translates into *in vivo* therapeutic value, we tested its efficacy to treat aggressive murine mixed-lineage-leukemia (MLL) eleven-nineteen leukemia (ENL)-driven p53WT acute myeloid leukemia (AML). Critically, MLL-ENL translocation has been shown to robustly increase Myc transcriptional programs [39]. Continued PMR-116 treatment at three different dosing regimens (72 mg/kg for 5 days; 120 mg/kg three times a week; 300 mg/kg once a week) notably delayed tumour progression monitored by bioluminescence imaging (**Figure 5A**). Consistent with the delay in tumour progression, overall survival of PMR-116 treated mice was increased by 206% in mice treated with 300 mg/kg (median survival vehicle: 15days, 72 mg/kg: 34 days, 120 mg/kg: 39 days, 300 mg/kg: 46 days; n=5 per group).

**Figure 5.**
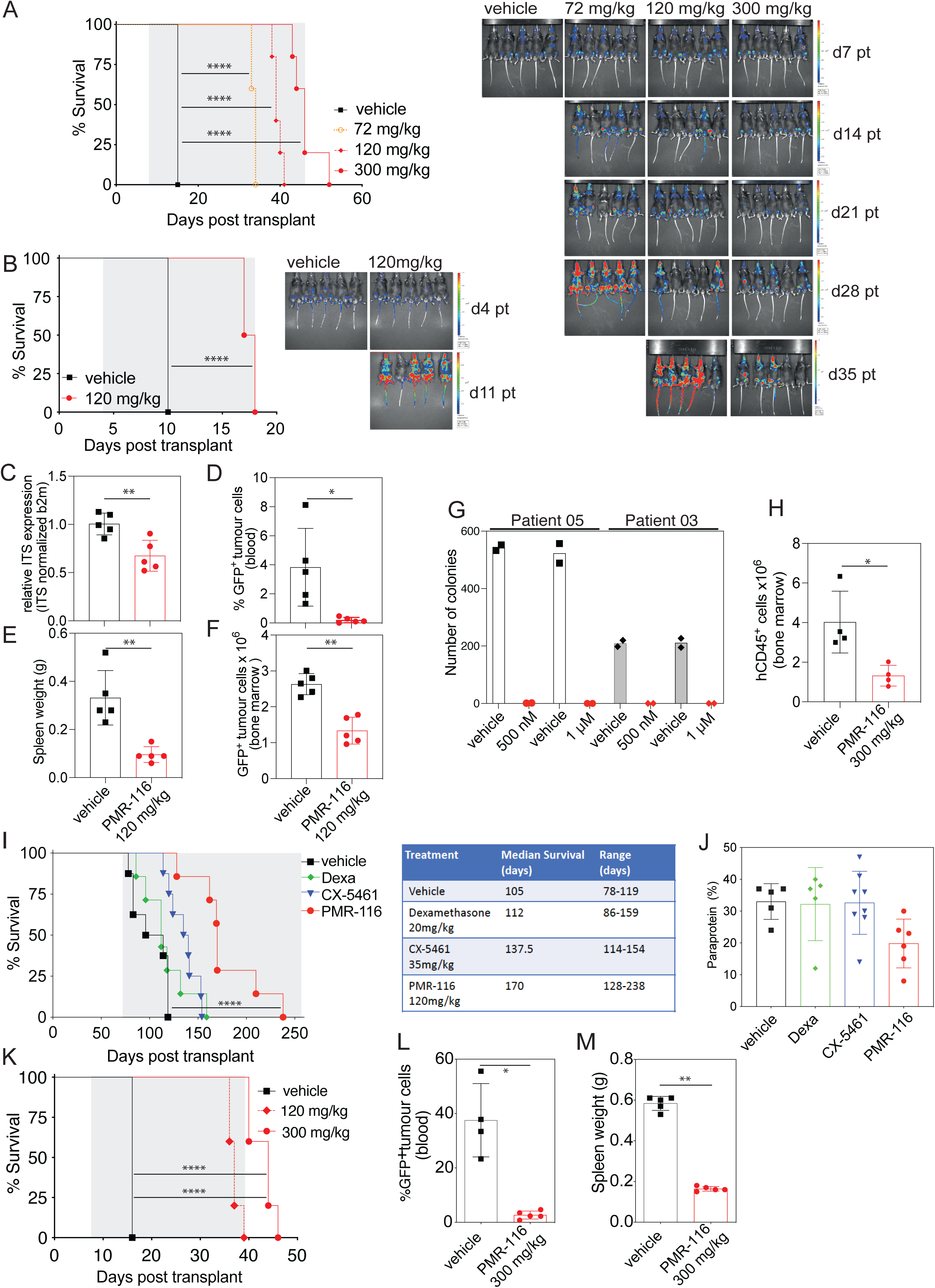
PMR-116 therapy treats aggressive haematological cancers. **A** PMR-116 therapy extends survival and delays tumour progression in MLL/ENL Nras driven p53WT AML (n=5 per group, **** *P* <0.0001 Log-rank test). **B** Survival is prolonged in p53null AML bearing mice receiving PMR-116 at 120 mg/kg (n=5 per group, **** *P* <0.0001 Log-rank test). **C** Orally administrated PMR-116 inhibits Pol I transcription 3 hrs post treatment in vivo (n=5 per group, ** *P* =0.0055 unpaired t-test). **D** Single dose PMR-116 treatment significantly decreases tumour burden in the peripheral blood (n=5 per group, *= 0.0162 unpaired t-test), **E** spleen (n=5 per group, **= 0.0021 unpaired t-test) and **F** bone marrow (n=5 per group, ** *P* = 0.0066 unpaired t-test) after 72 hrs in disease-bearing mice. **G** PMR-116 decreases colony forming capacity of primary human AML and **H** treats primary human AML in vivo (n=4 per group, *= 0.0166 unpaired t-test. **I** PMR-116 (120 mg/kg) therapy prolongs survival in murine V*κ*-MYC Multiple Myeloma (n=7-8 per group, **** *P* <0.0001 Log-rank test) and **J** reduces paraprotein (ns, one-way ANOVA non-parametric). **K** PMR-116 therapy at 120 and 300 mg/kg prolongs survival in Myc-driven Burkitt-lymphoma bearing mice (n=5 per group, **** *P* <0.0001 Log-rank test). Single dose PMR-116 administration at 300 mg/kg eradicated circulating disease in the peripheral blood (n=4/5 per group, * *P* = 0.0159 unpaired t-test) **L** and reduces splenomegaly **M** (n=5 per group, \*\**P*= 0.0079 unpaired t-test) in diseased animals after 24 hrs. Column graphs represent mean +/- SD

Furthermore, PMR-116 treatment slowed disease advancement in p53null AML, which correlated again with an attenuated but significant extension in survival compared to that observed in p53WT AML (75% increase in median survival vehicle: 10 days, 120 mg/kg: 17.5 days, n=5 per group) (**Figure 5B**). This suggests that PMR-116 therapeutic response in this AML model is at least in part independent of functional p53, consistent with literature indicating the activation of p53-independent NSP following nucleolar stress [40, 41]. Significant *in vivo* on-target (rDNA transcription) inhibition was confirmed in tumour-bearing mice after a single dose of the drug (120 mg/kg) (**Figure 5C**), demonstrating that even a modest decrease in Pol I transcription is sufficient to induce a robust reduction in tumour growth and increase in survival. Single-dose PMR-116 therapy (120 mg/kg) significantly reduced circulating disease as well as tumour load in the spleen and bone marrow of diseased mice (**Figure 5D-F**).

To evaluate whether PMR-116’s anti-cancer activity in murine AML translates into near-human disease, PMR-116 was tested for its efficacy in colony formation assays and primary patient-derived xenografts (PDX). In *ex-vivo* colony formation assays 500 nM and 1µM of PMR-116 inhibited the colony-forming capacity of two primary human AML models (de novo AML and undifferentiated AML) (**Figure 5G**). In an AML-derived PDX, 6 weeks of PMR-116 therapy delayed disease progression and significantly reduced the number of human CD45^+^ cells in the bone marrow of NSG mice compared to vehicle-treated animals (n=4; mean 1.323×10^6^ cells SD ±1.55 vs 4.028×10^6^ cells SD ±0.5) (**Figure 5H**).

Given that the sensitivity of PMR-116 correlates with elevated MYC activity, we further evaluated PMR-116 efficacy in Myc-driven murine haematological malignancies. PMR-116 demonstrated potent *in vivo* anti-tumour activity in V*κ*\*MYC multiple myeloma-bearing mice as evidenced by a 59% extended survival (**Figure 5I**) (median survival vehicle: 116.5 days n=8, PMR-116: 185 days n=7 and reduced increase in paraprotein (mean: vehicle 33% SD ±5.61vs PMR-116 19.8% SD±7.68) (**Figure 5J)**. Importantly, PMR-116 therapy demonstrated therapeutic efficacy superior to both multiple myeloma standard-of-care drug dexamethasone (Dexa) and first-generation Pol I inhibitor CX-5461 (median survival Dexa 129 days n=7; CX-5461 152.5 days n=8). In addition, PMR-116 was well tolerated over the 16 week treatment period; after changing the PMR-116 dosing regime from three times a week to twice weekly, the initial dip in white blood cell (WBC) counts slowly recovered (2wks treatment 5.2+/-1.61 ×10^9^/L; 4 wks treatment 6.629 +/-1.87 ×10^9^/L) (**SFigure 4**).

We also assessed PMR-116 in a highly aggressive Myc-driven B-cell lymphoma (Eμ-MYC) model, which revealed significant therapeutic benefit, whereby treatment at 120 mg/kg twice a week and 300mg/kg once weekly (QIW) with PMR-116 significantly extended overall survival in animals with lymphoma by 131.25% and 175%, respectively (**Figure 5K**) (median survival vehicle: 16 days, 120 mg/kg: 37 days, 300 mg/kg: 44 days). Single dose PMR-116 (300 mg/kg) significantly reduced circulating disease by 93% (vehicle 37.55% SD ±13.47 vs PMR-116 2.70% SD ±1.47, p<0.05) and splenic tumour burden by 72% (vehicle 0.58g SD±0.03 vs PMR-116 0.16g SD 0.01, p<0.01) (**Figure 5L&M**)

In summary, these findings demonstrate that PMR-116 is a promising novel cancer therapeutic drug with the potential to treat aggressive murine and human haematological cancers, particularly those driven by or with elevated MYC.

### RNA polymerase I inhibition is of therapeutic benefit in pre-clinical models of solid tumours

To evaluate the potential of PMR-116’s to treat solid cancers of different origin, PMR-116 was tested in murine and patient-derived models of prostate cancer, and in a Diethylnitrosamine (DEN) induced murine model of liver cancer which genetically and phenotypically closely recapitulates human hepatocellular carcinoma (HCC) [42, 43] . In the Hi-Myc mouse model of locally invasive prostate adenocarcinoma (characterized by the fragmentation or loss of smooth muscle actin (SMA) layer enclosing prostatic ducts (**Figure 6A**), PMR-116 significantly lowered the incidence of invasive lesions by ∼84% (10% vs 64%) analysed by histopathological scoring of Haematoxylin and Eosin (H&E) and ɑ-SMA of prostate tissue sections (**Figure 6B**). Consistent with this observation, a significant decrease in Ki67 positive cells (mean of 24.3% to mean 10.4%) but no significant increase in Caspase 3 cleavage positive cells were observed in the Hi-Myc prostate cancer-bearing mice treated with PMR-116 (**Figure 6C&D**). In marked contrast, the same dosing regime of PMR-116 provided no therapeutic benefit in the PTEN^-/-^ mouse model of prostate adenocarcinoma, as indicated by the disruption of ductal architecture and the lack of reduction in the percentage of Ki67 positive cells and absence of caspase 3 cleavage (**Figure 6E-G**). These results highlight the superior sensitivity of Myc-driven/overexpressed cancers to Pol I inhibition.

**Figure 6:**
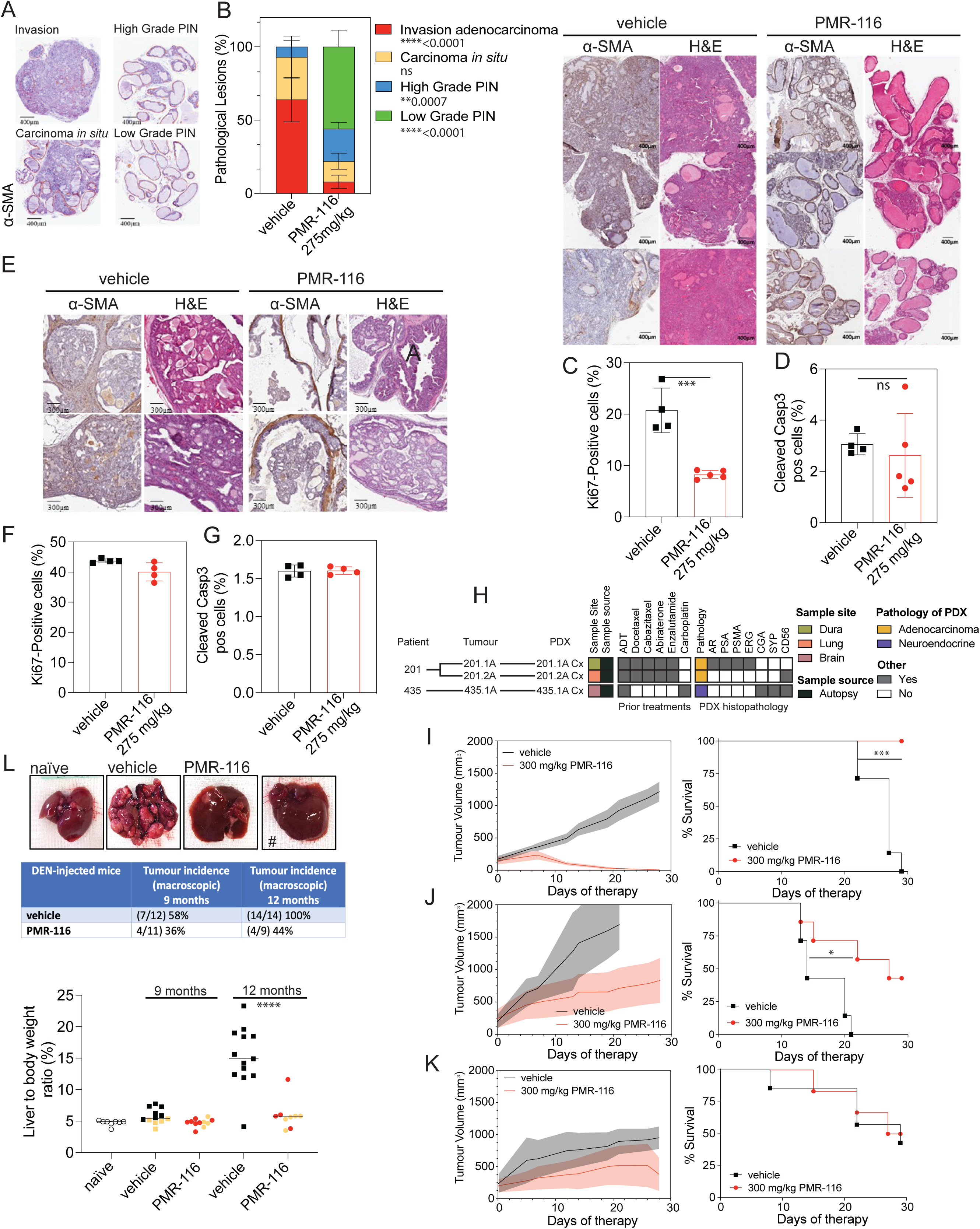
Selective Pol I inhibitor PMR-116 has therapeutic efficacy treating solid tumours *in vivo* **A** Representative images of α-SMA immunochemistry of 7 months old Hi-MYC to display the different phenotypical lesions within the model. **B** Continues PMR-116 treatment reduces the occurrence of invasive lesions in Hi-Myc prostate adenocarcinoma (n=5 per group, *** *P*=0.0007, ****<*P*=0.0001, unpaired t-test). Right panel: Representative images of α-SMA and Haematoxylin and Eosin H&E) immunochemistry of 7 month-old Hi-MYC mice orally treated with either PMR-116 (n=5, 275 mg/kg) or vehicle (n=4) (weekly, 5 doses) **C** PMR-116 treatment reduces the percentage of Ki67 positive cells (\*\*\**P*=0.0004, unpaired t-test) but does not significantly increase the percentage of cleaved Casp3 (CC3) positive cells **D** (ns, unpaired t-test) in the Hi-MYC lateral prostate of 7-month-old mice dosed with either vehicle (n=4) or PMR-116 (n=5) **E** Representative H&E and α-SMA stained images showing the pathological phenotype in anterior prostate of five-month-old PTEN-null mice after four weeks of treatment with PMR-116. **F** Quantitation of Ki-67 positive cells within the lateral prostate of five-month-old PTEN-null mice dosed with either vehicle (n=4) or PMR-116 (n=4) (weekly, 5 doses, ns unpaired *t*-test). **G** Quantitation of CC3 positive cells within the lateral prostate of five-month-old PTEN-null mice dosed with either vehicle (n=4) or PMR-116 (n=4) (weekly, 5 doses, ns, unpaired *t*-test). **H** Summary of the features of prostate cancer PDXs, including the patients’ previous treatments and the histopathology of the tumours. **I** Left: Changes in tumour volume (mm^3^) of PDX-435.1A-Cx in mice treated with vehicle (grey) or PMR-116 (orange; 300 mg/kg) for up to 27 days (n=7 per group). Solid line represents the average, shaded areas represent SD. Right: Kaplan-Meier survival curve showing median survival of 27 days for vehicle versus not reached for PMR-116 (\*\*\**P*=0.0002, Log-rank test). **J** Tumour growth and survival curve for PDX-201.1A-Cx (n=7 per group; \**P*=0.0139, Log-Rank test). **K** Tumour growth a survival curve for PDX-201.2A-Cx (n=7-6 per group; Log-Rank test, ns=non-significant). **L** PMR-116 exhibits potent in vivo antitumor activity in DEN-induced HCC. Representative liver images from 12-month-old DEN-injected mice treated once a week with either vehicle, CX-5461 (35 mg/kg) or PMR-116 (120 mg/kg). Control liver from DEN naïve mouse. Middle panel: Summary of macroscopic tumour incidence in DEN-injected mice after 9- and 12-month post injection. # indicates livers from treated DEN -injected animals with no detectable tumour. In number of mice with tumours in the liver and total number of animals per group). Lower panel: the graph presents the percentage of liver to body weight ratio in DEN-injected mice treated with vehicle or PMR-116 9 and 12 months post DEN injections (orange: no tumour, black: tumour vehicle, red: tumour PMR-116) (\*\*\*\**P*<0.0001, unpaired t-test). Control liver from drug and DEN naïve mice. Column graphs represent mean +/- SD.

PMR-116 was also tested in three PDXs of metastatic castration-resistant prostate cancer. These all originated from metastases of patients who died from prostate cancer after receiving standard-of-care therapies (**Figure 6H**). They represent androgen receptor-positive adenocarcinoma (201.1A-Cx), the double negative phenotype (201.2A-Cx) and small cell neuroendocrine prostate cancer (435.1A-Cx). PMR-116 treatment (300 mg/kg QW) significantly reduced tumour growth of all three PDXs (**Figure 6I-K, SFigure 6A-C & STable 1**). The effect was most striking for PDX 435.1A-Cx, with tumour regression in all mice in the PMR-116 treated group (**SFigure 6A**) and significantly longer survival; no mice reached the humane endpoint based on tumour burden (**Figure 6I**). PDXs 201.1A-Cx and 201.2A-Cx were established from different metastases of the same patient, and both had more moderate responses, with a reduced growth rate but no regression with PMR-116 treatment (**Figure 6 J&K, SFigure 6 B&C**). There was a significant increase in the survival of PMR-116 treated mice for 201.1A-Cx, but not 201.2A-Cx. In summary, PMR-116 suppresses the growth of diverse pathologies of metastatic castration-resistant prostate cancer that has progressed on currently available treatments.

We further expanded our study to test therapeutic efficacy of PMR-116 in the diethylnitrosamine (DEN) in vivo model of human hepatocellular carcinoma. Dysregulation of the c-Myc gene is a frequent occurrence in human HCC [44]. Myc plays a significant role in murine HCC induced by DEN [45]. PMR-116 demonstrated potent time-dependent *in vivo* anti-tumour activity this model of murine HCC at 3 and 6 months of therapy.

After 3 months of treatment, 7 out of 12 PMR-116 naïve/vehicle treated mice (58%) developed HCC. Remarkably, only 4 out of 11 PMR-116 treated mice (36%) developed HCC; tumours in these animals were smaller with an overall tumour growth inhibition (TGI) of 98% compared to HCCs in naïve mice (**Figure 6L**). After 6 months of treatment, all 13 mice (100%) in the PMR-116 naïve group harboured striking HCC burden with 12 out of 13 mice (92%) developing multiple, invasive large tumours. In contrast at 6 months, only 4 out of 9 PMR-116 treated mice (44%) developed HCCs; only 1 mouse (11%) developed tumours. Hence, an overall TGI of 89% with PMR-116 therapy (**Figure 6L**).

In summary, these findings demonstrate PMR-116 effectively and potently ameliorates growth of solid cancers in human- and relevant-murine models associated with elevated MYC.

### PMR-116 drug combination screen identifies drug combinations with improved anti-cancer activity

In clinical practice, single-agent therapy is rarely administered to cancer patients due to the increased risk of developing acquired resistance, causing disease relapse. To identify potential therapeutic agents that work synergistically with PMR-116 and further improve the therapeutic outcome and reduce toxicity in combination with PMR-116, a boutique small-molecule combination screen was performed in Mv4;11 cells. Drugs tested in combination with PMR-116 included FDA-approved agents and compounds in Phase I clinical trials for haematological cancers, including Acute lymphoblastic Leukemia, Chronic Lymphocytic Leukemia and AML, as well as a broad range of DNA damage, epigenetic and kinase inhibitors (**Figure 7A**). Drug combination synergy was assessed using the HSA and Bliss model of synergy [46, 47]. Cell viability analysis in the primary combination therapy screen revealed that out of the 316 therapeutic agents tested in combination with PMR-116, 10 drugs synergistically reduced cell viability at concentrations below 10 μM, including the kinase inhibitors amcasertib, arizotinib, dinaciclib, pazopanib, sunitinib, the cephalotaxine alkaloid homoharringtonine, the topoisomerase I inhibitor irinotecan, the PARP inhibitor niraparib, the anthracycline pirarubicin, and the aza-anthracenedione pixantrone maleate (**Figure 7B, SFigure 6**). The stemness kinase inhibitor amcasertib and the multi-targeted receptor tyrosine kinase inhibitor Sunitinib received the highest synergy score in both synergy models at concentrations below 10 μM (**Figure 7 & SFigure 7**). To confirm synergy or therapeutic antagonism in combination with PMR-116, small-molecule drugs that were identified as synergistic or antagonistic in the primary screen at concentrations below 10 μM were re-assessed in a secondary validation screen in Mv4;11 and KG-1 cells (**Figure 7C&D**). In total 33 compounds were re-assessed for their synergistic potential. Synergy scores >10 were confirmed in 14 out of 33 compounds, with the highest synergy observed for amcasertib, sunitib and the Bcl-2 inhibitor venetoclax in Mv4;11 (**Figure 7C**). In contrast, amcasertib showed little or antagonistic effect in KG-1 when combined with PMR-116, while the highest synergy with PMR-116 was achieved with Vinblastine in p53null KG-1 (**Figure 7D**). Overlaps in drug synergy (concentration < 10 μM)) in both cell lines (HSA score >10) was observed with the proteasome inhibitors carfilzomib and the second-generation HDAC1 inhibitor quisinostat (**Figure 7C&D**).

**Figure 7.**
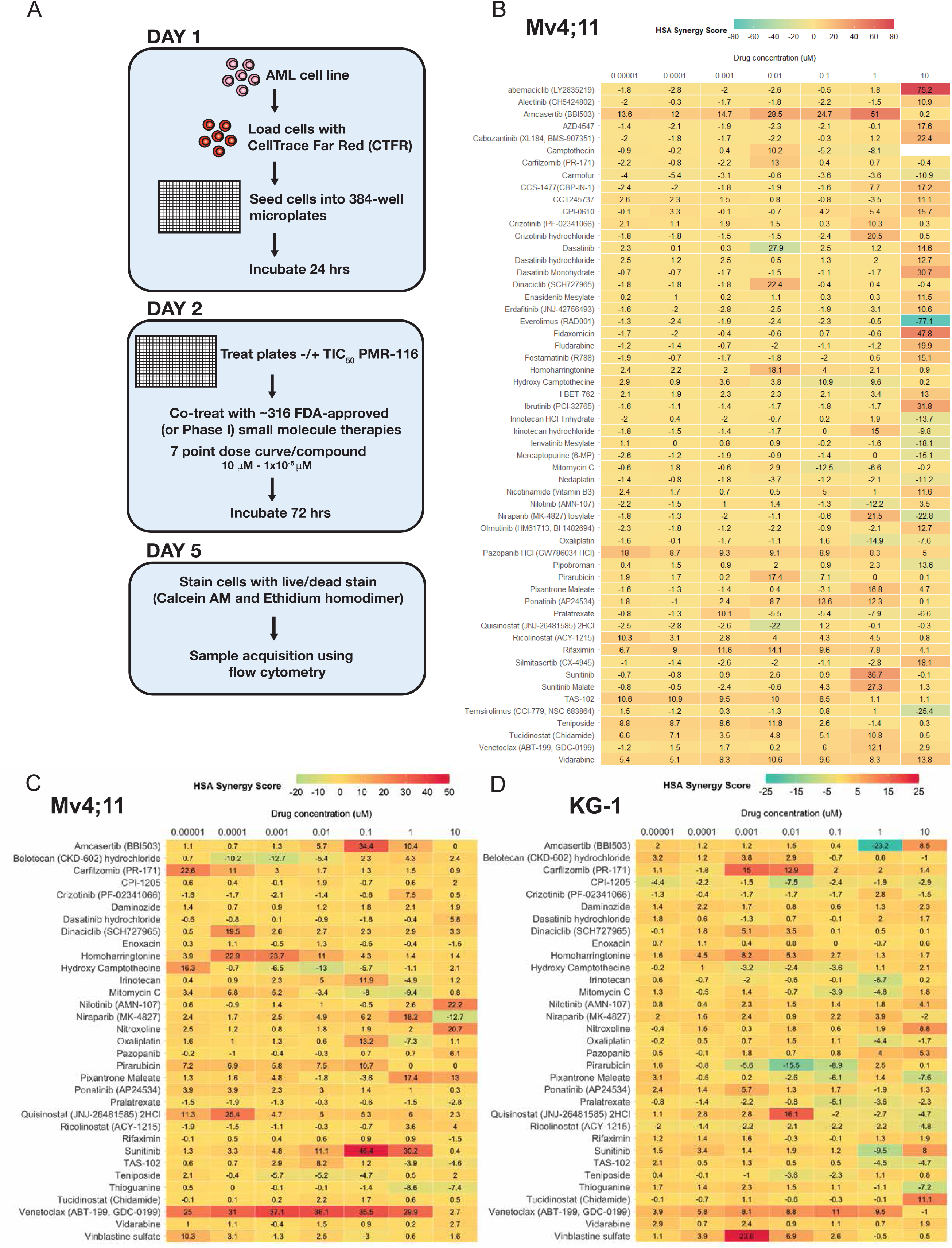
PMR-116 works synergistically with various clinically used compounds. **A** Schematic workflow of the primary drug combination screen in Mv4;11 cells. **B** Heat map of the primary combination screen displays the HSA synergy scores for each drug concentration (0.01nM, 0.1nM, 1nM, 10nM, 100nM, 1μM and 10μM) in combination with 400 nM PMR-116. Synergistic drug combination effects were assayed by changes in cell viability after 72 hrs of treatment. **C** and **D** HSA synergy score heat maps from the validation screen retesting compounds identified in the primary screen in Mv4;11 and KG-1 cells.

In conclusion, PMR-116 works synergistically, reducing cell viability, with several drugs used routinely for the treatment of haematologic malignancies in p53WT and null AML cell lines, and so providing possible new approaches for the treatment of haematologic malignancies with improved therapeutic outcome and reduce toxicity.

## Discussion

Since the initial discovery of CX-5461 a decade ago, the mechanism of action of this RNA Polymerase I transcription inhibitor remained in question, with several studies suggesting its efficacy is mediated through alternative mechanisms to inhibition of Pol I transcription and activation of the NSP, including direct poisoning of Top2⍺ to induce DNA damage and/or disruption of G quadruplex structures. Thus, whether directly targeting Pol transcription is a viable and efficacious approach to treat cancer has remained unclear. Here we report on the development of a second-generation Pol I transcription inhibitor, PMR-116, which exhibits improved drug-like properties and pharmacokinetics compared to CX-5461. Mechanistically, we demonstrate that on-target concentrations of PMR-116 (TIC90) inhibit rRNA synthesis most likely by stalling the Pol I complex at the rDNA promoter region and thus preventing promotor escape and resulting in reduced elongating Pol I. PMR-116 exhibits robust anti-cancer activity in a broad range of preclinical models of both haematologic malignancies and solid tumours, particularly those with elevated MYC. In contrast to CX-5461, the therapeutic response to PMR-116 occurs in the absence of global DNA damage suggesting that DNA damage is not an obligatory response to Pol I transcription inhibition but rather an off-target effect of the first-generation drug, most likely through the inhibition of Top2⍺ activity. Critically, our data establish that selective inhibition of RNA polymerase I is a *bone fide* therapeutic approach for cancer therapy and demonstrate that PMR-116 is a promising new drug with potential to treat a broad range of human solid and haematologic cancers where MYC is dysregulated and a driver of the malignancy.

Inhibition of Pol I transcription in response to certain extracellular stresses, e.g., radiation, chemotherapeutic drugs or genetic lesions in genes encoding the Pol I transcription apparatus, induce the NSP. NSP signalling is likely to have evolved to monitor nucleolar function and couple cellular proliferative capacity with RiBi activity to ensure they remain “hardwired” [38]. During NSP induction, inhibition of Pol I disrupts the equilibrium of rRNAs and ribosomal proteins (RPs) in the nucleolus, resulting in an accumulation of “free” RPs which exit the nucleolus and, through additional non-RiBi functions, activate tumour suppressors such as p53 and repress oncogenes such as MYC. The net effect is to block proliferation and induce cell cycle arrest, apoptosis and/or senescence [5, 16]. The best-described NSP (canonical NSP) involves the binding of RPL5 and RPL11 to the ubiquitin ligase MDM2, leading to the accumulation of the p53 tumour suppressor protein [48]. Consistent with PMR-116 inducing NSP, in p53 wt cells, PMR-116 induced rapid increases in p53 and p53 downstream signalling pathways. In addition to p53 signalling, transcriptome analysis revealed robust repression of MYC and E2F signalling, which are also well-described targets of the non-canonical NSP following inhibition of Pol I transcription and likely contribute to the therapeutic response of PMR-116. Consistent with non-canonical NSP pathways contributing to the therapeutic response, even though p53 null tumours were less sensitive to PMR-116 than p53 wt tumours, they nevertheless demonstrated a strong therapeutic response. Indeed, sensitivity to PMR-116 in a broad panel of cancer cell lines from solid tumours did not correlate with p53 status. Thus, we conclude that the therapeutic response to PMR-116 is likely mediated through the induction of a complex interplay between canonical and non-canonical NSP pathways.

As a single agent PMR-116 efficiently treats a broad range of pre-clinical models of both solid and haematologic tumours, including lymphoma, AML, hepatocellular carcinoma and prostate cancer, while sparing non-cancerous cells and tissues. Since RiBi is a ubiquitous cellular process, an important clinically relevant question to enable patient stratification in human trials is the understanding of the mechanism(s) that confer sensitivity or resistance to PMR-116. Our studies in normal immortalised cells and panel of cancer cell lines representing most major classes of malignancy clearly demonstrate that differential sensitivity between normal and tumour cells and between tumour cell types is not due to the differential ability to inhibit RNA Pol I transcription. Thus, while the process of malignant transformation significantly increases GIC50 sensitivity to PMR-116, the degree of sensitivity varies depending on the cancer cell type and even within cancers of similar origin. Our data support a mechanism in which the response to PMR-116 is dependent on how sensitive a given tumour type is to activation of the NSP and the subsequent downstream signalling to growth inhibition of apoptosis. This, in turn, is likely dependent on how the driver mutations affect ribosome biogenesis and nucleolar function. Thus, understanding which tumour cells will exhibit a strong NSP response is likely the key to predicting the therapeutic sensitivity of a given tumour to PMR-116.

Due to its primordial role in driving rDNA transcription and RiBi [11], high MYC expression seems to be a prime determinant of sensitivity to activation of the NSP in response to Pol I inhibition. We show that MYC expression positively correlates with sensitivity to PMR-116 in human-derived cancer cell lines **(Figure 2B)**. We also provide compelling evidence that high MYC tumours are particularly sensitive to Pol I transcription inhibition as evident by the significant delay in tumour progression in response to PMR-116 in mice bearing V*κ*\*-MYC multiple myeloma, Em-Myc B-cell lymphoma, high MYC PC and also HCC and MLL-ENL driven AML (both which exhibit high MYC levels) **(Figure 5&6)** [39, 44, 45]. Moreover, we provide some evidence that it is the driver-mutation rather than cancer type that determines response to Pol I inhibition, supported by the observation that PTEN^-/-^-driven prostate adenocarcinoma was completely resistant to PMR-116 compared to MYC-driven prostate adenocarcinoma, which was exquisitely sensitive. These results were somewhat surprising given that PTEN negatively regulates phosphoinositide 3-kinase (PI3K)/Akt/mammalian target of rapamycin (mTOR) signalling pathways to control proliferative cell growth and survival in part through repression of Pol I transcription by interfering with SL-1 complex formation [49]. Thus, not all driver mutations that directly modulate RiBi sensitise cells to Pol I therapies. The centrality of *MYC* in oncogenesis makes it a highly attractive target, but despite advances in understanding *MYC*, targeting it directly has been largely unsuccessful due to its intrinsically disordered nature, precluding structure-guided drug design [50]. Our data provide strong evidence that targeting RiBi with PMR-116 exploits a critical therapeutic vulnerability in high *MYC* cancers, suggesting that PMR-116 has clinical promise to treat a broad range of human solid and haematologic cancers where MYC is dysregulated and a driver of the malignancy.

Using a focussed molecule drug combination screen in AML we demonstrated that PMR-116 acts synergistically in reducing cell viability with of the 10 drugs tested, including the kinase inhibitors amcasertib and sunitib, and the BCL2 inhibitor venetoclax, all used routinely for the treatment of haematologic malignancies. Of particular interest was venetoclax, which showed strong synergy with PMR-116 at low doses in p53WT Mv4;11 AML, but less so in p53null KG1, consistent with intact p53 function being essential for sustaining durable responses to BH3-mimetic drugs like venetoclax [51]. This synergy is consistent with how these two drugs might cooperate to regulate the p53 apoptosis axis where PMR-116 induces stabilisation of p53 through activation of the NSP and regulates apoptosis upstream of BCL-2, whereas venetoclax drives apoptosis downstream by inhibiting negative regulators like BCL2 [52].

Based on the above data and a favourable toxicology profile, we have gained IND approval and launched a Phase I clinical trial of PMR-116 in patients with advanced solid tumours (CTRN12620001146987). The toxicity, response, pharmacokinetic and co-relative data from this trial will offer valuable insights as to whether selective inhibition of RiBi independent of DNA damage reported with CX-5461, might offer a new non-genotoxic therapeutic approach in the treatment of cancers, particularly those driven by MYC, which are difficult to treat.

## Material and methods

### Tissue culture

Cell lines were either purchased from DSMZ or ATCC. Mv4;11 and KG-1 cells were maintained in RPMI-1640 supplemented with 20% heat inactivated Fetal Bovine Serum (FBS) and GlutaMax (Thermo). PC3 and LNCap were maintained in RPMI 1640 supplemented with 10% FBS. Cell lines included in the growth inhibitory studies were cultured in either Eagle’s Minimum Essential Medium, Dulbecco’s Modified Eagle’s Medium, RPMI-1640 Medium, F-12K Medium, IMDM, FBS and Antibiotic-Antimycotic (Gibco) according to ATCC and DSMZ instructions.

### Transcription inhibitory concentration (TIC)50 determination

Cell lines were seeded 24 (Mv4;11, KG-1) or 48 hrs (PC3, LNCaP) prior to drug treatment. Cells were treated with 3, 10, 30, 100, 300, 600, 1000, 3000, and 10 000 nM of PMR-116 or vehicle (50 mM citric acid) for 3 hrs and total RNA was extracted using a Nucleospin RNA kit (Macherey-Nagel, cat. 740955.250) according to manufacturer’s instructions. cDNA was prepared using a SuperScript IV Reverse Transcriptase kit (Invitrogen). Quantitative polymerase chain reactions (qPCR) were performed on the StepOne Plus Real-Time PCR System (Applied Biosystems) using a SYBRGreen Fast Master Mix (Applied Biosystems) and primers specific for 5’ETS, cFOS, MYC and the housekeeping gene B2M (primer sequences see STable 2). For each biological replicate, expression was normalised B2M via the delta Ct method [2-(*Δ*Ct)]. Expression changes were further normalised to the vehicle (treatment/vehicle) and the dose response curve obtained with GraphPad Prism 10 using non-linear regression ([inhibitor] versus response – variable slope (four parameters)) and the target IC50 (TIC50) calculated.

### In vitro transcription assays

Non-specific transcription assays were performed as described previously [53]. Specific transcription reactions were performed as described in [33, 54] Supercoiled plasmid DNA (prHu3) or linear DNA fragment (Fr440, free or mobilized on the beads) containing the human rRNA gene promoter were used as a template. Reactions were supplemented with either HeLa Nuclear extract (NE) or with appropriate purified factors [33, 53, 54] S1 nuclease protection was performed as described in [54] with a 5’-end labelled oligonucleotide identical to the region between –20 and +40 of the template strand in the human rRNA gene promoter. The amount of RNA produced by *in vitro* transcription reactions was quantified using a phosphorimager, FLA-7000 (Fuji) and Aida software.

### Analysis of Pol I PIC

HeLa nuclear extract (NE) or purified factors were incubated with the immobilized DNA template (IT) as previously described [54] and PIC formation was evaluated by *in vitro* transcription assay and western blot analysis. Proteins were eluted in 8 M urea buffer and analysed by Western blotting with antibodies specific to human Pol-I subunit A190, UBF, and SL1 subunit TAF1B (see STable 1).

### Transcriptomics analysis

Mv4;11 and KG-1 cells were treated with either 3xTIC50 (1 µM) PMR-116, 1 µM CX-5461 or their corresponding vehicle (50 mM citric acid, 50 mM NaH_2_PO_4_) for 3 and 12 hrs. Total RNA was extracted using Nucleospin RNA kit (Macherey-Nagel 740955.250) per manufacturer’s instructions. RNA library preparation and sequencing were performed at the Biomolecular Resource Facility (BRF) at ANU. RNA library preparation was performed using the Illumina stranded mRNA prep kit and pooled RNA library were sequenced on the NovaSeq6000 S1 flowcell, 200 cycles, 2×100 bp paired end. The RNA-seq analysis pipeline was implemented in R, gene-level counts were extracted using tximport after mapping with Salmon (v1.5.20 and the human reference genome (GRCh38.p13 from GENCODE (v37). The pipeline included quality control, time course differential expression analysis using DESeq2, and gene enrichment analysis.

### Chromatin Immunoprecipitation (ChIP)-seq

Mv4;11 cells, PMR-116 (1 µM), CX-5461 (1µM) or vehicle (50 mM citric acid and 50 mM NaH_2_PO_4_) for 30 min, were cross-linked with 0.8% formaldehyde and chromatin was immunoprecipited with anti-Pol I (RPA127) or anti-UBF antibodies (STable 3) as described in [17] and DNA was purified using ChIP DNA clean and concentrator kit (Zymo Research D5205) according to the manufacturer’s guide.

Sequencing libraries were prepared using NEBNext Ultra II DNA Library Prep Kit for Illumina from New England BioLabs and sequenced on the NovaSeq 6000 (S1flow cell 100 cycles, 2 x 50bp paired end. For the bioinformatic analysis a custom human reference genome was created as follows to facilitate the measurement of ChIP-seq signal across rDNA. Briefly, rDNA reference sequence (GenBank Accession No. KY962518.1) was fragmented in-silico to generate overlapping 150 bp sequences with a step size of 1bp. The generated pseudo-reads were aligned to the human reference genome (GRCh38.p13), allowing up to 1000 possible matches, using Bowtie2 version 2.3.5.1 [55]. The regions across the genome that were identified as ‘rDNA-like’ were masked using ‘maskfasta’ tool in BEDTools suite version 2.30.0 [56], and a single rDNA repeat sequence was incorporated as an extra chromosome. The sequencing reads were aligned to the custom reference using Bowtie2 with default parameters. Normalised coverage tracks for ChIPed samples were generated using the tool bamCompare in deepTools suite version 3.4.3 [57]. The resulting coverage tracks were used for visualisation through Integrative Genomics Viewer (IGV) [58]

### Growth inhibitory concentration (GIC) 50 determination

Cells (3*10^4^ cells/well) were plated in 96-wells plates and incubated overnight (adherent) or treated directly (suspension) in triplicates with PMR-116 (14 nM, 41 nM, 124 nM, 370 nM, 1.11 μM, 3.33 μM, 10 μM). Following 96 hrs incubation Alamar Blue Cell Viability assay (Invitrogen DAL1100) was performed per manufacturer instructions. Fluorescence (ex 545; 590 nm) was measured with SPECTRAmax® M5 microtiter plate reader (Molecular Devices). Fluorescence values were background-corrected (subtraction of average blank controls) and 100% viability was assigned using the average of untreated controls (UTC). Percentage viability was expressed as a percentage of UTC. These values were fitted against logarithms of corresponding PMR-116 molar concentrations with GraphPad Prism 10 to non-linear regression, sigmoidal dose-response (variable slope) model using appropriate constrains (e.g. bottom must be between 0.0 and 5.0) to determine corresponding GIC50. In case of cell lines, for which multiple experiments were carried out, median, mean and standard error of means values were calculated with Column Statistics analysis.

MYC expression data were received from publicly available RNA seq data set at the EMBL-EBI Expression Atlas, plotted as transcripts per million, n=1-5 per cell line.

### Cell death and cell cycle analysis

Cells were seeded approximately 24 hrs prior treatment with 300 nM – 3 μM PMR-116 or CX-5461 or the corresponding concentration of vehicle (50 mM Citric Acid, 50 mM NaH_2_PO_4_). Cells were stained at 24, 48 and 72 hrs post treatment using APC Annexin V Apoptosis Detection Kit with 7-AAD (BioLegend, 640920) as per manufacturer’s instructions and analysed using BD LSR II cytometer. Acquired data were analysed using FlowJo^TM^ v10.8 Software (BD Life Sciences). Cells were analysed at 12 and 24 hrs post-treatment with vehicle, PMR-116 or CX-5461. An hour prior to harvest, cells were pulse-labelled with 10 μM BrdU. After harvesting, cells were fixed with ice cold 80% ethanol, treated with 2N HCl/ 0.5% (v/v) Triton ×100 and neutralized with 0.1M Na_2_B_4_O_7_. BrdU incorporation was probed with anti-BrdU antibody and anti-IgG-A488 (see Table S3) and total DNA was stained with 10 μg/ml 7AAD.

### Immunoblotting

Cells were lysed in Western Solubilisation Buffer (20mM HEPES, 0.5mM EDTA, 2% SDS, 0.1M DTT and protein concentration was determined by using DC Protein Assay (BioRad 5000116) as per manufacturer’s instructions. Proteins were separated by SDS-polyacrylamide gel electrophoresis and transferred to Immobilon-P PVDF membrane (Millipore, IPVH00010) using Trans-Blot Turbo Transfer System (BioRad). The membranes incubated with primary antibodies overnight at 4°C and with secondary antibodies for 1 hr at RT. Protein bands were visualised by using Clarity Western ECL Substrate (BioRad 1705061) in the ChemiDoc System (BioRad) with Image Lab software (version 6.0.1, 2017).

### *γ*H2AX immunofluorescence

Mv4;11 and KG-1, seeded at 3×10^5^ cells/mL, and treated with vehicle, 3x TIC50 CX-5461 or PMR-116 for 24 hrs and 5 μM Doxorubicin (Selleckchem S1208) for 3 hrs were cytospun onto Poly-L-lysine coated glass slides (Sigma-Aldrich, P0425). PC3 and LNCaP were grown for 48 hrs on glass coverslips and treated with 3x TIC50 of CX-5461 or PMR-116 or vehicle for 24 hrs or 5µM Doxorubicin for 3 hrs. Cells were fixed with 4% Paraformaldehyde (PFA; Electron Microscopy Sciences, 15714S), permeabilized with 0.5% Triton X-, blocked with PBS/5%BSA, prior to incubation with anti-*γ*H2AX (at 4°C) and anti-IgG-A647 (Life Tech, A21235: at 37°C). After staining with 5 μg/mL DAPI in PBS (Sigma, D-9542), slides were mounted using Fluorescence Mounting Media (Agilent, S3023) and imaged using LSM800 confocal microscope (Zeiss) with a 63X (1.4) lens. Images were analysed using Fiji v2.14. For each treatment condition, DAPI^+^ nuclei were selected and the mean fluorescence intensity (MFI) of A647 per nuclei was calculated. A minimum of 3 fields of view and 15 cells were analysed per treatment condition. The average MFI of 3 independent biological replicates per treatment condition was analysed using Tukey’s multiple comparison test (GraphPad Prism 10).

### Nucleolar Morphology and G4 quadruplex immunofluorescence

Mv4;11, KG-1, PC3 and LNCaP cells were seeded and treated as described above. After 6 hrs cells were fixed with 2% PFA. For suspension cells 2.25×105 cells per treatment were cytospun onto glass slides. Cells were permeabilized with 0.3% TritonX-100 and incubated 1 hr at 37°C with anti-Fibrillarin and anti-Nucleolin and secondary (IgG-A594, AgG-A647) antibodies. Slides were stained with DAPI and mounted as described above.

For G4 stabilisation analysis cells were treated with 3x TIC50 of PMR-116 and 10 µM of TMPy4 and corresponding vehicles (50 mM Citric acid or ultra-pure H_2_O) for 1 and 3 hrs. Cells were pelleted, fixed and permeabilized as described above and treated with 40 μg/mL RNase for 1 hr at 37°C, incubated 1h at 37°C with anti-G4 and 1h, RT with anti-IgG-A647 antibody. Analysis was performed as mentioned above.

### AML Patient colony cultures

Colony formation capacity of primary patient AML cells was analysed in methylcellulose (Stemcell Technologies M4435) as described previously [16]. Patient consent was obtained under human ethics number ETHLR16.141.

### PMR-116 in vivo efficacy testing in preclinical cancer models

All animal experimentations were performed according to protocols approved by the Peter MacCallum and Australian National University Animal Ethics Committees. An ethical endpoint was reached when mice show enlarged lymph nodes, early hindlimb paralysis, 20% weight loss, or general debility such as hunched, ruffled, or reduced mobility.

### Pharmacokinetic (PK) analysis

PK analysis of plasma and brain mouse samples was conducted by Medicilon Preclinical Research (Shanghai) LLC. A single dose of 100 mg/kg PMR-116 was administered via oral gavage to male ICR mice. Blood samples from three animals per timepoint were collected via cardiac puncture at 15 min, 30 min, 1, 2, 4, 8, 12 and 24 hrs post dose. Blood samples were placed into tubes containing sodium heparin and centrifuged conditions at 8000 rpm for 6 minutes at 4°C to separate plasma from the samples. Following centrifugation, the resulting plasma were transferred to clean tubes and stored frozen at -80°C pending bioanalysis. The brain of each animal was harvested, excised and rinsed by saline, dried on filter paper and snap frozen on dry ice and then stored at -80°C until further analysis. The concentrations of PMR116 in plasma and brain homogenate samples were determined using a high-performance liquid chromatography/mass spectrometry (HPLC/MS/MS) method. The PK parameters from mean concentration-time data were calculated using a non-compartmental module of WinNonlin® Professional 5.2. Any BLQs (LLOQ = 1.0 ng/mL for plasma; LLOQ = 5.0 ng/g for brain) were omitted when calculate the PK parameters.

### AML transplant models

AML *in vivo* experiments were conducted under approved animal ethics protocols (A2018/48 and A2021/38) at ANU. C57/Bl/6 male mice were transplanted with 5 x 10^5^ MLL/ENL Nras (p53 WT or p53null) AML cells/mouse via tail vein injection. Seven days post-transplant, the mice were randomized into four groups and dosed continuously via oral gavage with either vehicle (PBS, 50 mM citric acid) or PMR-116 at 72 mg/kg QD (5 days a week), 120 mg/kg (3 days a week) or 300 mg/kg (once a week) until an ethical endpoint was reached. Engraftment and disease progression were monitored once a week by bioluminescence imaging using the IV/VIS Xenogen after injection of mice with luciferin (200 μL per mouse of a 1 mg/ml stock solution). Mice were weight at least three times a week. For single dose short term therapy experiments mice were transplanted as described above and treated when disease was established with a single dose of vehicle or PMR-116 (300 mg/kg) via oral gavage and tissue obtained for subsequent analysis at the indicated timepoints.

### AML Patient-Derived Xenotransplantation (PDX)

*In vivo* experiments were conducted under approved animal ANU ethics protocols (A2018/48 and A2021/38). Male and female NSG mice (n=9) were irradiated once with 1Gy one day prior to transplant. Mice were transplanted intravenously with 10^6^ primary patient derived AML cells via tail vein injections. Animals were randomized into treatment group 6 months post-transplant when circulation disease was detectable in the peripheral blood. Mice were treated with either 300 mg/kg PMR-116 or vehicle (50 mM citric acid) for 6 weeks via oral gavage, then mice were harvested and the number of human CD45^+^ cells within the bone marrow was analysed by flow cytometry.

### Multiple Myeloma

Experiments were performed in accordance with approved animal ethics protocol E462 at the Peter MacCallum Cancer Centre, Australia. C57BL/6 mice were irradiated 2x with 3Gy 6 hrs apart using a Caesium source. The next day 5×10^5^ V*κ*\*MYC cells (4929 clone provided by Prof. Ricky Johnstone at Peter MacCallum Cancer Centre) were injected per mouse. Paraprotein levels were determined using standard serum protein electrophoresis (SPEP) by PeterMacCallum Pathology. Mice were randomized into 4 groups at the time of detectable level of paraprotein and treated with vehicle control, 35 mg/kg CX-5461, 20 mg/kg dexamethasone and 120 mg/kg PMR-116. PMR-116 therapy commenced at 3 times a week which was changed to twice weekly at the end of week 2 due to weight loss. CX-5461 and dexamethasone was given once a week orally. Mice were monitored twice weekly, and serial bloods were used to monitor tumour burden via SPEP.

### B-Cell lymphoma

*In vivo* lymphoma experiments were performed at ANU under approved animal ethics projects (A2015/12, A2018/48). Mice were i.v. injected with 2×10^5^ Eµ-MYC GFP cells (4242 clone) and engraftment was confirmed 9 days post-transplant by flow cytometric analysis of GFP+ cells in peripheral blood. Mice were randomized into groups and treated with either 120 mg/kg PMR-116 3x per week, 300 mg/kg PMR-116 1X per week or vehicle (50 mM citric acid) via oral gavage. For short term therapy experiments mice were transplanted as described above and treated when disease was established with a single dose of vehicle or PMR-116 (300 mg/kg) via oral gavage. Tissue was collected for subsequent analysis at the indicated timepoints. At harvest, mice were anaesthetized and a cardiac bleed performed. Blood was processed for WBCC (Advia) and flowcytometric analysis of B220/GFP positive cells in the blood. Spleen weights were recorded.

### Murine Prostate Cancer

Murine prostate cancer experiments were conducted under the approved animal ethics protocol E581 at the PeterMac Callum Cancer centre. Hi-MYC (ARR2pbsn-MYC) and PTEN-deficient (Pb-Cre^+/-^PTEN^fl/fl^) mice were obtained from breeding colonies at Monash University, Australia [18]. Hi-MYC and PTEN-deficient male mice were aged between 6–7 months and given a weekly dose of 275 mg/kg PMR-116 by oral gavage for four weeks to assess drug efficacy over a chronic regime. For acute experiments, each animal was given a single dose of PMR-116 (275 mg/kg) 24 hrs prior to analysis. Prostate lobes were harvested with 1/2 (lateral prostate, dorsal prostate, ventral prostate, anterior prostate) preserved in formalin for Immunohistochemistry (IHC).

### Histopathology & Immunohistochemistry (IHC)

Lateral prostate lobes were harvested, formalin-fixed for 48 hrs, paraffin embedded and 5 μm thick sections with prepared. No prostate sections were discarded due to the heterogeneity of PCa within the prostate. Every 20^th^ section was stained with haematoxylin & eosin (H&E) (Dako) to visualise lesion formations and histopathological tumour progression. These sections were given a percentage based on the different tumour pathological features displayed by the prostatic lobes. These scores were attributed by at least two researchers who were blind to the genotype and treatment group. Fragmentation of normal prostate architecture was measured using smooth muscle actin, which in most cases surrounds epithelial ducts in tumour-free mice. For Hi-MYC mice, each duct was given a percentage score for each type of lesions: Low/High grade PIN, carcinoma *in situ* (CIS), and invasive adenocarcinoma. Specifically, this data was expressed as the percentage of ducts with one of these lesions out of the total number of ducts in the analysed sections. As PTEN null mice do not present with invasive adenocarcinoma, their lesions were scored based on the percentage of CIS present in the analysed section.

Specific antigens were quantified using immunohistochemistry. Relevant sections underwent dewaxing and antigen retrieval Citrate with (pH6) or Tris-EDTA (pH9) and were then incubated with Hydrogen Peroxide (Dako) and CAS blocking reagents (Dako). Sections were then incubated with a specific primary antibody antibody diluent (Dako) for 1h at RT followed bt goat α-mouse or goat α-rabbit immunoglobulins in Dako Envision (Dako EnVision®+ Dual Link System-HRP (DAB+), K4065) for 1h, at RT. Antigen visualisation was done by briefly incubation in chromogen D-amino benzidene (DAB) (Dako). Certain antibodies requiring specific isotype secondary were incubated with Avidin-Biotin Complex (ABC) (Thermofisher) to amplify antigen signal. DAB was then used to reveal specific staining. Sections were counterstained in Haematoxylin for 30 secs, mounted and scanned digitally using an Aperio AT turbo slide scanner (Leica Biosystems). Positively stained cells were quantified with algorithms built into the Aperio Image Scope Analysis software (Leica Biosystem).

### Prostate cancer PDXs

Serially transplantable prostate cancer PDXs were previously established by the Melbourne Urological Research Alliance (MURAL)[59] with informed, written consent from patients in accordance with human ethics approvals from the Peter MacCallum Cancer Centre (11/102) and Monash University (12287). The PDXs in this study were established from samples collected by the CASCADE program at the Peter MacCallum Cancer Centre [60]. All experiments were performed according to Monash University animal ethics approvals (28911, 22555, 30132, 36038). PDXs were routinely authenticated with STR profiling and histopathology review, including confirming the absence of lymphoma. For treatment experiments, PDX tissues were regrafted subcutaneously into pre-castrated, male, immune-deficient NSG mice. Once grafts reached approximately 100 mm^3^, mice were systematically allocated to the vehicle (50 mM citric acid) or PMR-116 (300 mg/kg) group and treated weekly by oral gavage. Graft volumes were measured three times per week with callipers. Mice were humanely killed if the grafts reached 1000 mm^3^. The remaining animals were given their final dose of treatment 24-hour prior to being humanely killed. Changes in tumour volume were analysed with linear mixed models as previously described [61].

### Diethylnitrosamine (DEN) induced liver cancer model

In vivo therapy experiments in liver cancer bearing animals were approved under the ANU animal ethics A2017/13. C57Bl/6 male mice purchased from APF and 12-15 days old mice were injected intraperitoneally with either saline (n=5) or 10 mg/kg Diethylnitrosamine (DEN, N0756 Sigma-Aldrich; n=60). At 6 months after DEN injection disease progression was confirmed and DEN injected animals were randomized into treatment groups (120 mg/kg PMR-116 once a week, 35mg/kg CX-5461 twice a week and vehicle) dosed by oral gavage.

At 9 and 12 months post DEN injection and 3 and 6 months of treatment a cohort of vehicle, PMR-116 and CX-5461 treated animals were harvested, therefore mice were weighed, anaesthetized; blood obtained by cardiac puncture. Full blood counts were obtain using the Advia. Once the liver was removed and weighed, it was cut into 3 sections: 1 section was diced into small pieces and split over 2 tubes, one for RNA and the other for protein extraction. 1 section was placed in formalin and the last diced and placed in a cryo-mould in OCT, frozen on dry ice, covered in foil and stored at -80°C for IHC. A full necropsy was performed to detect any additional lesions. Tumour burden within the liver was assessed by determining the liver to body weight ratios (LBWR) of naïve strain-gender- and age-match mice from LBWR of mice HCC. Tumour growth inhibition (TGI) was calculated using the following equation: TGI (%) = [1(aveLBWR_PMR116_/aveLBWR_Vehicle_)] *100.

### Combination drug screen

A 316-compound library of small molecules containing FDA-approved and Phase I compounds, as well as CDK, DNA Damage, Epigenetic and Kinase Inhibitors used to treat haematological malignancies and myeloproliferative disorders was purchased from SelleckChem (see Stable 3 for compounds and catalogue numbers).

Prior to seeding, cells were labelled using the CellTrace Far Red Proliferation Kit (ThermoFisher 34564) as per the manufacturers’ protocol and resuspended in culture media. Cells were seeded into 384 well clear tissue culture plates (Corning 3701) at a density of 8000 cells/well and incubated for 24 hours in a humidified microplate incubator (Liconic) at 37°C with 5% CO_2_. Microplates were treated with vehicle or 1x TIC50 dose of PMR-116 and simultaneously with a 7-point dose curve of the aforementioned compounds (for most drugs, 10 μM, 1 μM, 100 nM, 10 nM, 1 nM, 0.1 nM and 0.01 nM; for those with limited solubility, the highest treatment dose was 2 μM) prepared in cell culture media delivered into microplates using the JANUS G3 robot (PerkinElmer). Microplates were then incubated for 72 hrs in a humidified microplate incubator at 37°C with 5% CO_2_. Controls on all plates included DMSO (negative control) and dinaciclib (100 nM final concentration, positive control) with and without PMR-116 treatment.

Culture media was removed using the JANUS G3 robot, and cells were stained with a mixture of 10nM calcein-AM and 1uM ethidium homodimer-1 from LIVE/DEAD Viability/Cytotoxicity Kit (ThermoFisher L3224) and CountBright absolute counting beads (ThermoFisher C36950). Microplates were incubated for 20 minutes at 37°C with 5% CO_2_ and analysed with iQue Screener Plus (Sartorius) with a 10 second sip, 2 seconds up (speed = 29 rpm) using ForCyte software (Sartorius), collecting data outputs including the % Live Cells, % Dead Cells and Median CTFR signal intensity parameters. For each assay plate, quality control (QC) analysis data was collected and Z-factor (Z’) were calculated comparing the negative and positive controls across the % Live cells and Median CTFR signal [62] (average -/+ 2SD -0.89 -/+ 0.15 and 0.80 -/+ 0.10, respectively).

Live Cells and Median CTFR intensity) and SynergyFinder 2.0 [63] was used to calculate synergy, an open source software available at median executed using R .

## Acknowledgements

We would like to thank the Cytometry, Histology and Spatial Multiomics (CHaSM) Facility (Dr. Harpreet Vohra and Mr. Michael Devoy), the staff at the Australian Phenomics Facility in particular Anthony Baker and the Centre for Advanced Microscopy (Dr Daryl Webb, Dr Angus Rae and A/Prof Melanie Rug).

Further we would like to thank the staff at the Australian Phenomics Facility, the ACRF Biomolecular Resource Facility (BRF) and the ANU Centre for Therapeutic Discovery (ACTD; in particular, Dr Jinshu He, Ms Kate Parsons, Dr Maxim Nekrasov at the John Curtin School of Medical Research at the Australian National University for their assistance.

This work was supported by funding from the National Health and Medical Research Council (NHMRC) of Australia (Project Grants #1158732, Ideas Grant #2002741 and Senior Research/Investigator Fellowships to R.D.H (#1116999, #2009504). Funding for the combination therapeutic screening study was also supported by a Maddie Riewoldt’s Vision (https://mrv.org.au; grant-in-aid; #MRV18) and Therapeutic Innovation Australia (NCRIS) pipeline voucher awarded to A.J.G and R.D.H.

The BRF and ACTD (A.J.G) were established through Australian Cancer Research Foundation funding (ACRF) in partnership with the Australian National University, and both receive operational funding from the Australian Government’s National Collaborative Research Infrastructure Strategy (NCRIS) program through Phenomics Australia (ACTD), Therapeutic Innovation Australia (ACTD) and Bioplatforms Australia (BRF).

K. Maclachlan received research funding from the Leukemia and Lymphoma Society.

## Authors contribution

R.F., K.M.H., D.D., L.F., R.D.H. and N.H. conceived, designed and supervised the study.

R.D.H. and N.H. wrote the manuscript; all other authors have reviewed and edited the manuscript. R.D.H., J. H. and A.J.G. designed and performed the combination drug screening utilized in this study. Preclinical studies were conducted and analysed by N.H., K.M.H., K.M., R.R., A.H., S.H., M.L., G. R., R.T., A.C., G.F., S.S., N.T.. *In vitro* experiments were performed and analysed by R.F, K.M.H., K. P., P.P., M.D., V.B., E.R., M.A., P. N., M. G. B., Y. M., D.D. and N.H.. The computational analysis was done by T.U., A.C., Z.Y. and E.E.. M.H. designed and provided PMR compounds.

## Supplementary figure legends

**SFigure 1.**
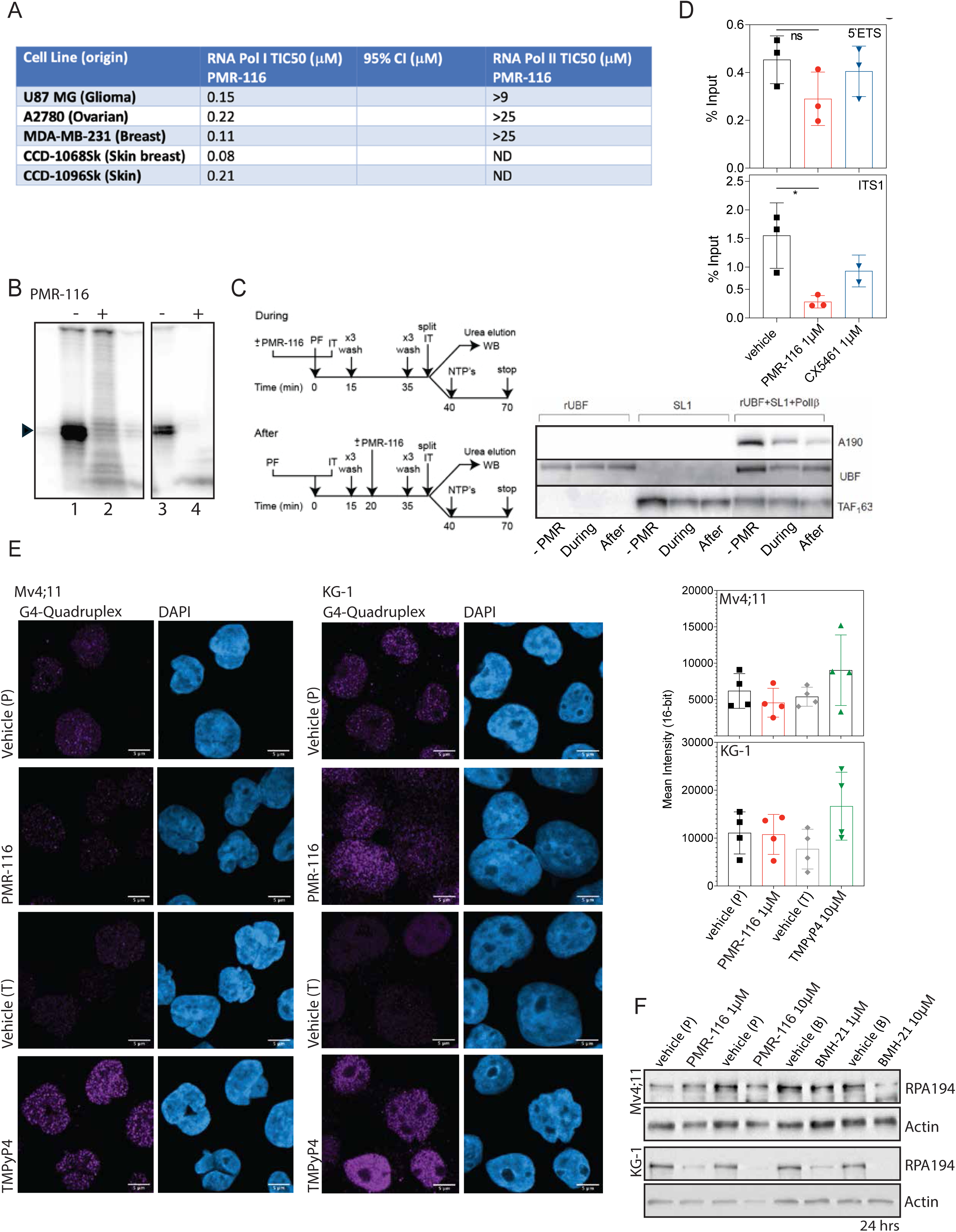
PMR-116 selectively inhibits RNA polymerase I in the absence of interference of PIC formation and G4 stabilisation. **A** Summary table of RNA Pol I and Pol II TIC50 for PMR-116 in human glioma, ovarian, breast cancer and normal cells. **B** Specific transcription reactions containing 2 µl of HeLa NE and 100 ng of supercoiled plasmid DNA template (left panel) or 2 µl Do2, 1 µl SL1 and 100 µg of linear template (right panel) were supplemented with 100 µM of PMR-116 as indicated. Transcripts were analyzed by S1 nuclease protection assay; a representative image is shown. Protected radiolabelled fragments, reflecting accurately initiated transcripts of 40 nt, are indicated by an arrowhead. Lanes 1 and 3 are the control reaction containing no drug. **C** Schematic presentation of the experimental workflow (left panel) PMR-116 (50 µM) was added during (Dur) or after (Aft) PIC assembly with purified factors on an immobilized rDNA template (IT). ND is a no drug control. For immunoblotting membranes were probed with antibodies against largest subunit of Pol I A190, UBF, and SL1 subunit TAF1B. Representative image from two independent experiments is shown (right panel). PMR-116 has no detectable effect on the binding of UBF, but causes a notable decrease in TAF_1_63 binding and reduces the amount of UBF and Pol I A190 bound to the template in the presence of SL1 when drug is present during and after PIC formation. **D** PMR-116 reduces PAF53 association with the transcribed region of the rDNA. Graphs present PAF53 occupancy as percentage input at 5’ETS and ITS1 rDNA region in Mv4;11 cells treated with either vehicle, PMR-116 (1µM) or CX-5461 (1µM) for 30 min (ns = non-significant, \*\**P*=0.0026 (1µM), one-way ANOVA Dunn’s multiple comparison). n=3 independent biological replicates **E** Representative images of DNA quadruplex structure (G4) immunofluorescence in Mv4;11 and KG-1 cells treated with PMR-116 (1µM), TMPyP4 (10µM) or the respective vehicle for 3 hrs, in contrast to TMPyP4 PMR-116 does not stabilizes G4 structures. DAPI (blue) marks the nucleus, G4 structures (pink). Scale bars represent 5µm, magnification 63x. Right panel: Quantification of G4 immunofluorescence presented as the mean intensity of 4 independent biological replicates **F** Immunoblot analysis of Pol I large subunit RPA194 and Actin in Mv4;11 cells treated with vehicle, 1uM PMR-116 or BMH-21 for 6 hrs. Representative image from a minimum of n=3 independent biological replicates.

**SFigure 2.**
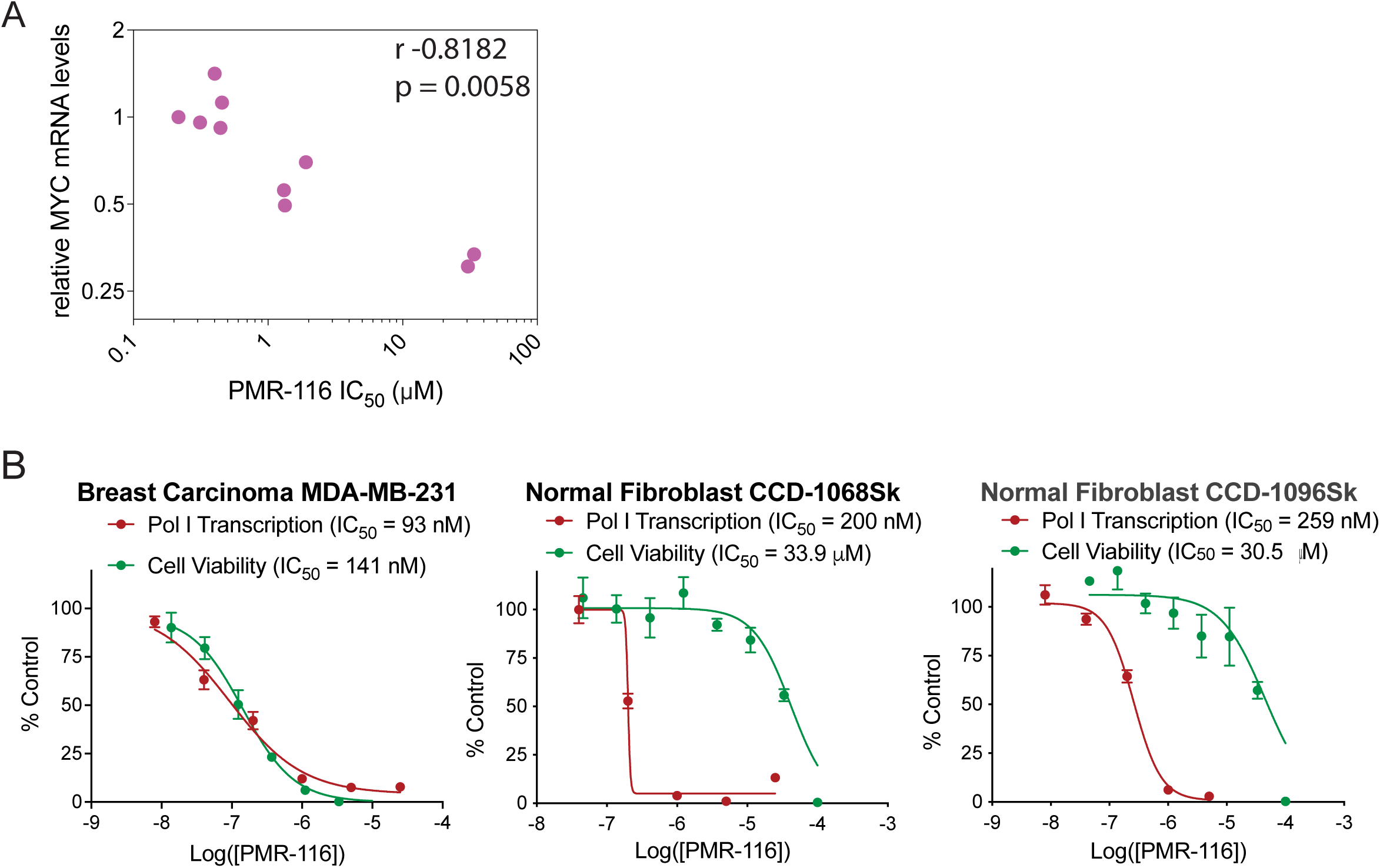
Sensitivity to PMR-116 correlates with MYC expression. **A** Correlation plot displays PMR-116 IC_50_ and relative MYC mRNA expression analysed by qPCR in triple negative breast cancer (TNBC). Spearman correlation r= -0.08182; *\*\* P*= 0.0058. **B** PMR-116 has similar on target activity in cancer (breast carcinoma: MDA-MB-231) and normal cells (normal fibroblasts: CCD-1068Sk, CCD-1096Sk) with PMR-116 more potently reducing cell viability in cancer cells compared to normal cells compared to the control. Relative transcription and cell viability are plotted as % of the respective control. Graph represents at least n = 3 independent biological replicates.

**SFigure 3.**
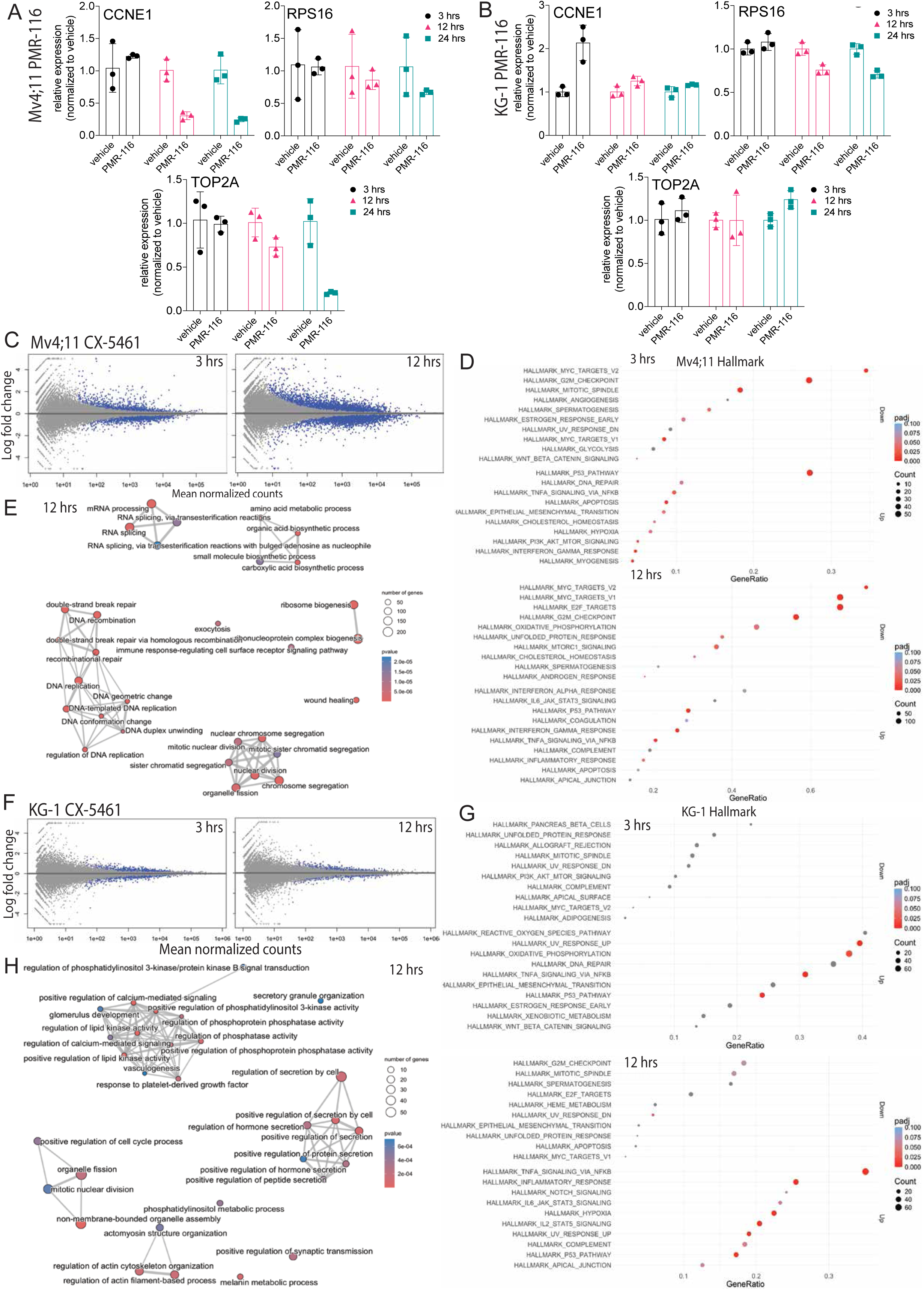
Pol I inhibition induces differential gene expression in Mv4;11 and KG-1 cells. **A** and **B** validation of differential expressed genes (CCNE1, RPS16, TOP2A) identified in the RNA seq in Mv4;11 and KG-1 cells after 3, 12 and 24 hrs of PMR-116 treatment via qPCR. Graph represents the mean +/- SD of n = 3 independent biological replicates. **C** and **F** M (log ratio) A (mean average) plots display all differentially up or downregulated genes in response to CX-5461 treatment after 3 and 12 hrs in Mv4;11 and KG-1. Blue dots indicate significant differentially transcribed genes compared to vehicle. **D** and **G** Hallmark pathway analysis at 3 and 12 hrs post CX-5461 treatment in Mv4;11 and KG-1. Fast gene set enrichment analysis with pre-ranked geneset by log2FC*(-log10(pvalue)) and a minimum number of 10 genes per pathway and a maximum of 500 genes size per pathway. **E** and **H** Enrichment Map analysis of the top 30 deregulated pathways in both Mv4;11 and KG-1 after 12 hrs of CX-5461 treatment.

**SFigure 4.**
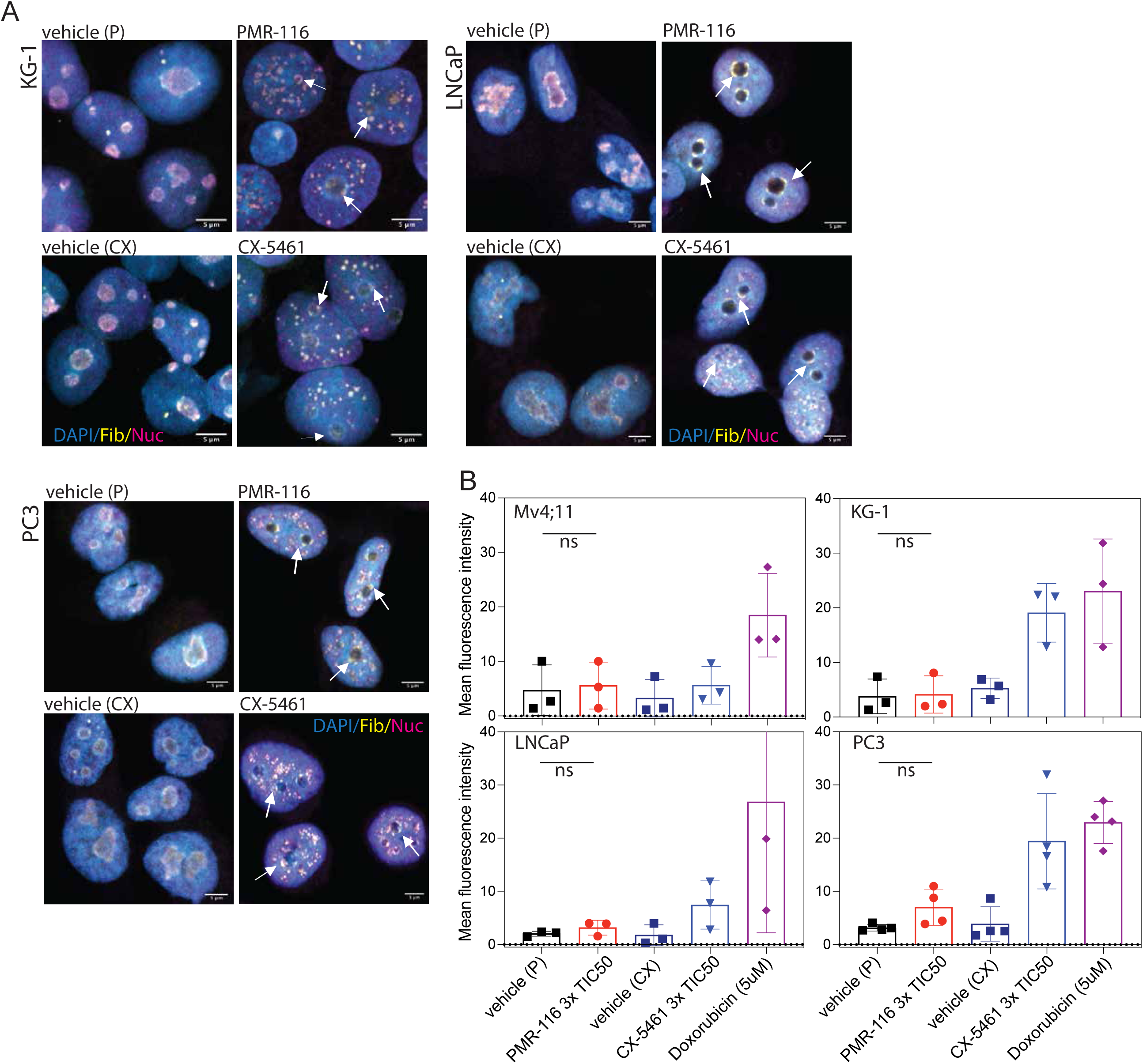
PMR-116 treatment induced nucleolar rearrangement in haematological and prostate cancer cell lines. **A** Changes in nucleolar morphology was assessed by Fibrillarin (yellow) and Nucleolin (pink) immunofluorescent staining in KG-1, PC3 and LNCap cells treated for 3 hrs with either vehicle (citric acid or NaH_2_PO_4_), PMR-116 (3xTIC50) or CX-5461 (3xTIC50). Arrows indicate nucleolar cap formation in response to treatment. CX-5461 treated cells served as a positive control. The nucleus was stained using DAPI (blue). Images were taken at 63x magnification, scale bar represent 5µm. Represented are representative images of n = 3 independent biological replicates. B Quantification of the IF *γ*H2AX foci IF. Presented as the mean fluorescence of n=3-4 independent biological replicates. Ordinary one-way ANOVA Tukey’s multiple comparison test. ns=non-significant

**SFigure 5.**
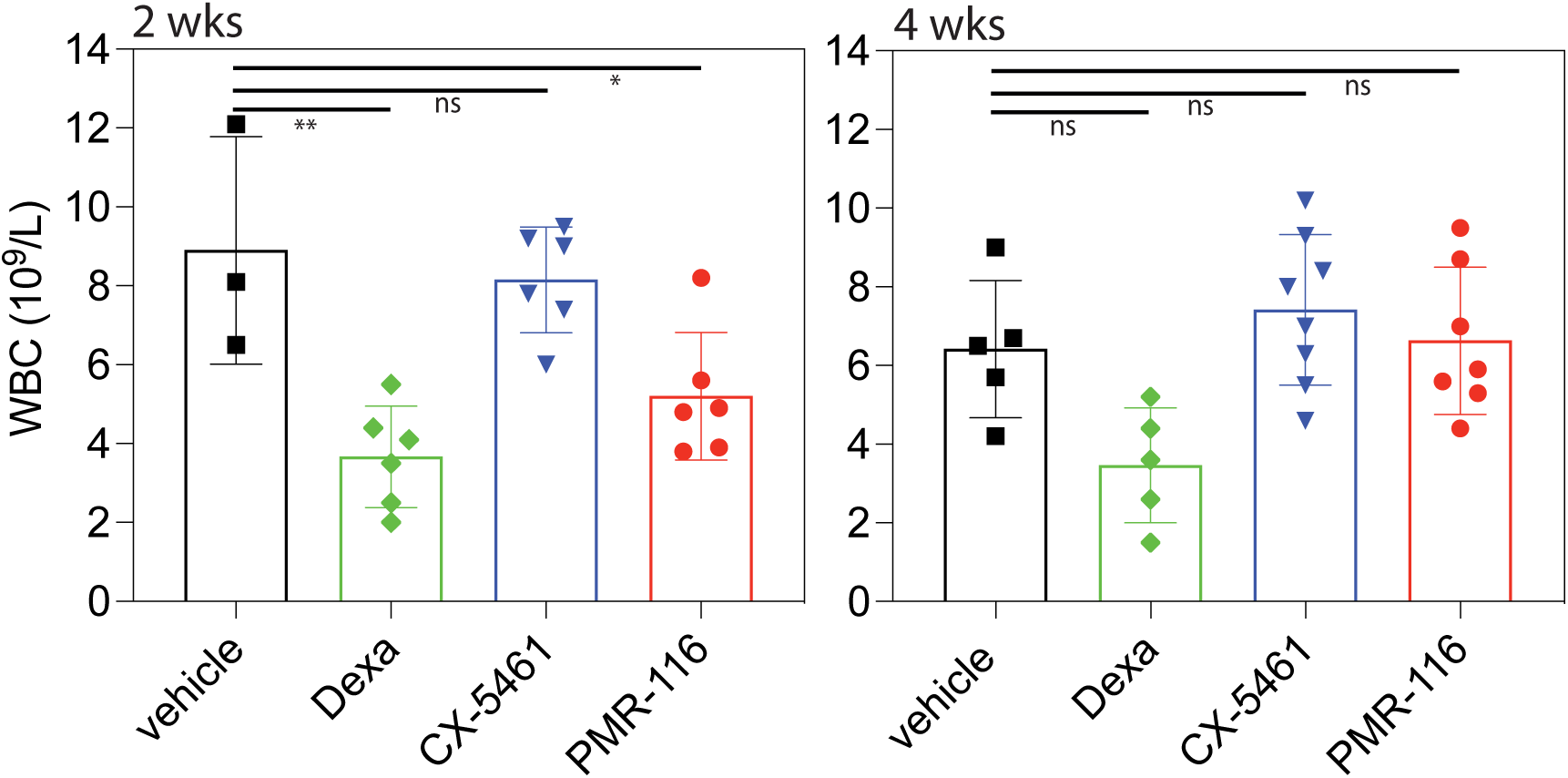
WBC analysis in V*κ*-Myc multiple myeloma bearing mice after 2 wks of therapy, treated with either vehicle, dexamethasone (Dexa) 20 mg/kg, CX-5461 35 mg/kg or PMR-116 and after 4 wks of treatment when the dosing regime was reduced from three times weekly 120 mg/kg to twice weekly. n= 3-8 per treatment group. Ordinary one-way ANOVA Tukey’s multiple comparison test, \*\**P*=0.0018, \**P*=0.0271, ns=non-significant.

**SFigure 6.**
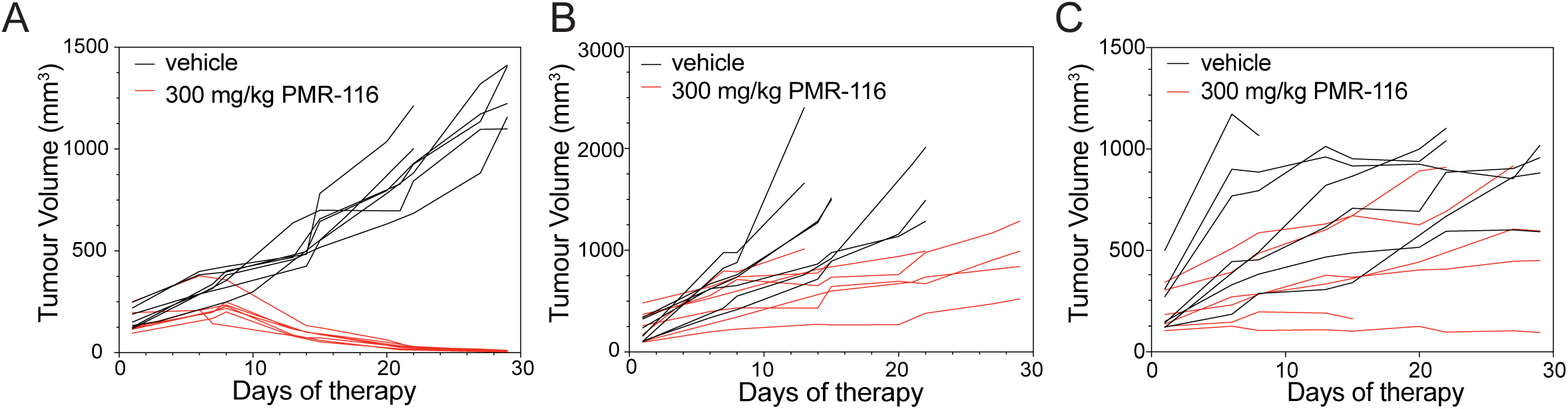
Multidrug resistance metastatic prostate cancer primary derived xenografts display varied sensitivity to PMR-116 therapy. Individual tumour volume for 435.1A-Cx (**A**), 201.1A-Cx (**B**) and 201.2A-Cx (**C**) derived prostate PDX treated with either vehicle or 300 mg/kg PMR-116 (n=7-6 per group).

**SFigure 7.**
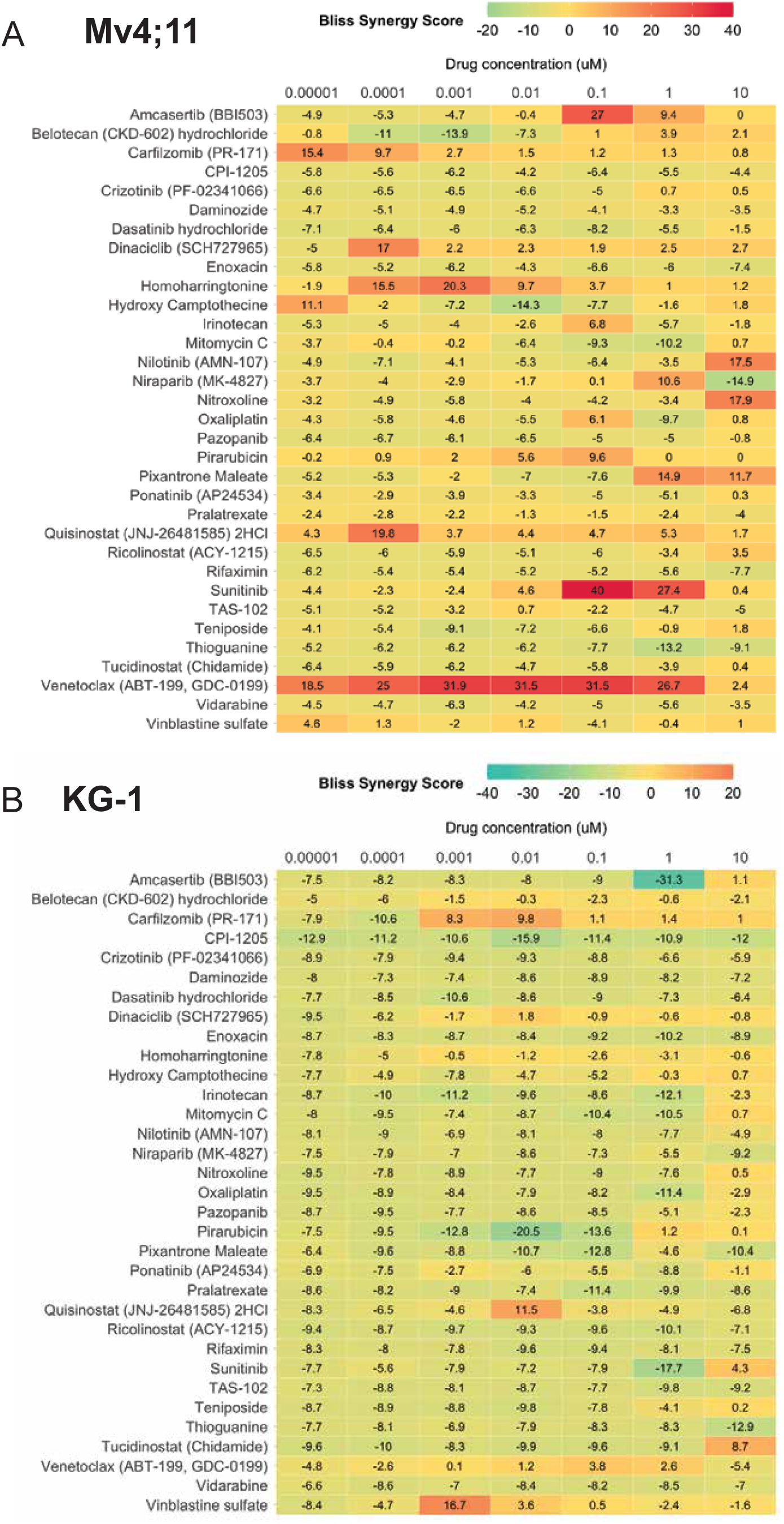
PMR-116 has synergistic therapeutic efficacy in Mv4;11 and KG-1 cells. Heat maps of validation screens in Mv4;11 (**A**) and KG-1 (**B**) display the Bliss synergy scores for each drug concentration tested (0.01nM, 0.1nM, 1nM, 10nM, 100nM, 1μM and 10μM) in combination with 400 nM PMR-116. Synergistic drug combination effects were assayed by changes in cell viability after 72 hrs of treatment.

**Table S1.** Linear mixed model analysis of tumor volume following treatment with PMR-116.

| Treatment | Treatment vs vehicle<br>*P* value (95% confidence intervals) |
| --- | --- |
| **201.1A-Cx** |  |
| PMR-116 (300 mg/kg) | 0.002 (-722.5 - -199.1) |
| **201.2A-Cx** |  |
| PMR-116 (275 mg/kg) | 0.150 (-467.5 - 77.6) |
| PMR-116 (300 mg/kg) | 0.027 (-608.7 - -41.0) |
| **435.1A-Cx** |  |
| PMR-116 (300 mg/kg) | <0.001 (308.8 - 571.5) |

**Statistical analysis of graft volume within each treatment group across time**
| Treatment | Tumor volume reached significance compared to day 0 of treatment | *P* value (95% confidence intervals) | *P* value (95% confidence intervals) | *P* value (95% confidence intervals) | *P* value (95% confidence intervals) | *P* value (95% confidence intervals) | *P* value (95% confidence intervals) | *P* value (95% confidence intervals) | *P* value (95% confidence intervals) |
| --- | --- | --- | --- | --- | --- | --- | --- | --- | --- |
| **201.1A-Cx** |  |  |  |  |  |  |  |  |  |
| **Day** |  | **5** | **7** | **13** | **14** | **19** | **21** | **26** | **28** |
| Vehicle | Day 5 (significantly increased) | <0.001 (-642.8 - -245.2) | <0.001 (-712.6 - -315.1) | <0.001 (-1282.6 - -855.1) | <0.001 (1304.6 - -887.5) | <0.001 (-1529.5 - -1112.4) | <0.001 (-1637.9 - -1220.8) | Vehicle harvested | Vehicle harvested |
| PMR-116 (300 mg/kg) | Day 5 (significantly increased) | 0.028 (-394.5 - -22.7) | 0.006 (-466.9 - -80.0) | <0.001 (524.23 - -152.4) | <0.001 (-594.1 - -222.2) | <0.001 (-631.7 - -244.6) | <0.001 (-688.4 - -301.2) | <0.001 (-803.6 - -397.5) | <0.001 (-849.7 - -443.6) |
| **201.2A-Cx** |  |  |  |  |  |  |  |  |  |
| **Day** |  | **5** | **7** | **12** | **14** | **19** | **21** | **26** | **28** |
| Vehicle | Day 5 (significantly increased) | <0.001 (-487.2 - -247.4) | <0.001 (-512.1 - -272.3) | <0.001 (-638.6 - -398.8) | <0.001 (-651.3 - -411.5) | <0.001 (-704.9 - -465.1) | <0.001 (-782.1 - -542.3) | <0.001 (-807.3 - -567.5) | <0.001 (-839.9 - -600.1) |
| PMR-116 (275 mg/kg) | Day 5 (significantly increased) | <0.001 (-369.2 - -129.4) | <0.001 (-394.2 - -154.4) | <0.001 (-456.6 - -216.8) | <0.001 (-478.0 - -238.2) | <0.001 (-468.34 - -228.5) | <0.001 (-532.9 - -293.1) | <0.001 (-555.5 - -269.3) | <0.001 (-606.8 - -340.7) |
| PMR-116 (300 mg/kg) | Day 12 (significantly increased) | 0.225 (-209.222 - 49.8) | 0.060 (-253.6 - -5.4) | 0.009 (-303.6 - -44.6) | 0.004 (-319.2 - -60.192) | <0.001 (-406.8 - -132.8) | <0.001 (-428.1 - -154.1) | <0.001 (-489.6 - -195.4) | <0.001 (-452.7 - -128.1) |
| **435.1A-Cx** |  |  |  |  |  |  |  |  |  |
| **Day** |  | **6** | **7** | **12** | **14** | **19** | **21** | **26** | **28** |
| Vehicle | Day 6 (significantly increased) | <0.001 (-239.8 - -83.4) | <0.001 (-266.2 - -116.7) | <0.001 (-394.0 - -237.6) | <0.001 (-534.4 - -384.9) | <0.001 (-694.3 - 537.9) | <0.001 (-830.6 - -681.0) | <0.001 (-1022.5 - -872.9) | <0.001 (-1121.0 - -971.4) |
| PMR-116 (300 mg/kg) | Day 6 (significantly decreased); Day 18 (significantly increased) | 0.041 (-152.0 - -3.4) | 0.023 (-161.9 - -12.3) | 0.233 (-29.6 - 119.9) | 0.100 (-12.146 - 137.4) | 0.004 (36.3 - 185.9) | 0.009 (33.416 - 230.1) | <0.001 (59.9 - 209.4) | <0.001 (64.4 - 214.0) |

**Statistical analysis of graft volume compared to vehicle control across time**
| Treatment | Treatment tumor volume reached significance compared to vehicle control | *P* value (95% confidence intervals) | *P* value (95% confidence intervals) | *P* value (95% confidence intervals) | *P* value (95% confidence intervals) | *P* value (95% confidence intervals) | *P* value (95% confidence intervals) | *P* value (95% confidence intervals) | *P* value (95% confidence intervals) |
| --- | --- | --- | --- | --- | --- | --- | --- | --- | --- |
| **201.1A-Cx** |  |  |  |  |  |  |  |  |  |
| **Day** |  | **5** | **7** | **13** | **14** | **19** | **21** | **26** | **28** |
| PMR-116 (300 mg/kg) | Day 13 | 0.232 (-126.3 - 499.5) | 0.227 (-125.3 - -508.6) | <0.001 (383.9 - 1009.7) | <0.001 (320.5 - 957.7) | <0.001 (511.3 - 1156.8) | <0.001 (563.0 - 1208.5) | Vehicle harvested | Vehicle harvested |
| **201.2A-Cx** |  |  |  |  |  |  |  |  |  |
| **Day** |  | **5** | **7** | **12** | **14** | **19** | **21** | **26** | **28** |
| PMR-116 (275 mg/kg) | Not reached | 0.332 (-151.5 - 340.5) | 0.333 (-151.6 - 430.4) | 0.162 (-87.4 - 494.6) | 0.180 (-96.1 - 485.8) | 0.080 (-32.9 - 549.1) | 0.067 (-561.8 - 20.2) | 0.053 (-3.471 - 596.6) | 0.074 (-27.8 - 564.9) |
| PMR-116 (300 mg/kg) | Day 5 | 0.040 (16.1 - 621.8) | 0.052 (-3.367 - 602.4) | 0.017 (73.2 - 678.9) | 0.018 (70.2 - 676.0) | 0.028 (40.86 - 652.2) | 0.012 (96.8 - 708.3) | 0.019 (66.5 - 686.1) | 0.006 (144.5 - 777.5) |
| **435.1A-Cx** |  |  |  |  |  |  |  |  |  |
| **Day** |  | **6** | **7** | **12** | **14** | **19** | **21** | **26** | **28** |
| PMR-116 (300 mg/kg) | Day 6 | 0.025 (13.883 - 200.0) | 0.006 (37.6 - 218.2) | <0.001 (291.4 - 477.6) | <0.001 (455.5 - 636.1) | <0.001 (657.7 - 843.8) | <0.001 (800.9 - 1021.4) | <0.001 (1015/6 - 1196.2) | <0.001 (1118.7 - 1299.3) |

**Table S2.**
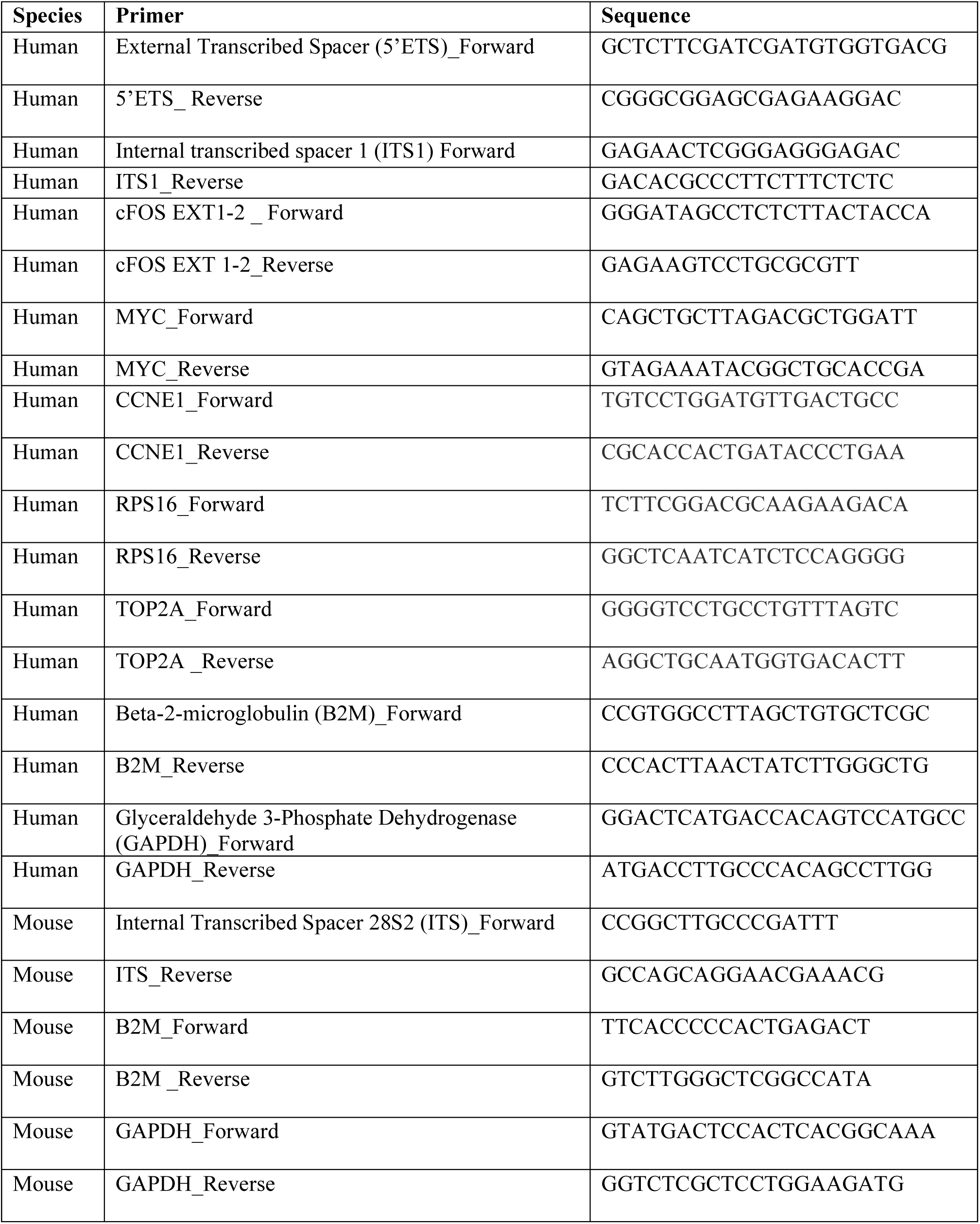

**Table S3.**

**STable 3**
| **Antibody** | **Catalogue** | **Application** | **Dilution/volume** |
| --- | --- | --- | --- |
| Chk1 (2G1D5) | Cell Signaling 2360S | WB | 1:1000 |
| Chk1 | AB Clonal A4194 | WB | 1:1000 |
| Chk2 | Cell Signaling 2662S | WB | 1:1000 |
| Chk2 | AB Clonal A19543 | WB | 1:1000 |
| Phospho-Chk1 (S345) (133D3) | Cell Signaling 2348S | WB | 1:500 |
| Phospho-Chk1 (S345) | AB Clonal A21009 | WB | 1:1000 |
| Phospho-Chk2 (T68) (C13C1) | Cell Signaling 2197S | WB | 1:500 |
| p21 waf1/cip1 (12D1) | Cell Signaling 2947S | WB | 1:1000 |
| p53 (DO-1) | Santa Cruz SC-126 | WB | 1:2000 |
| Phospho-p53 (S15) | Cell Signaling 9284S | WB | 1:1000 |
| RPA194 | Santa Cruz Sc-48385 | WB | 1:1000 |
| RPA194 (F-6) | Santa Cruz SC46699 | WB | 1:250 |
| Rabbit anti TAFI63 sera | In-house | WB | 1:1000 |
| βActin (C4) – HRP conjugate | Santa Cruz SC-47778 | WB | 1:10,000 |
| Goat Anti-Rabbit IgG (H+L) – HRP conjugate | BioRad 1706515 | WB | 1:2000 |
| Goat Anti-Mouse IgG (H+L) – HRP conjugate | BioRad 1706516 | WB | 1:2000 |
| Rabbit anti mouse/human Fibrillarin | Abcam ab5821 | IF | 1:250 |
| Mouse anti human Nucleolin | Abcam ab136649 | IF | 1:250 |
| Anti-DNA G-quadruplex (G4) antibody, clone 1H6 | Sigma-Aldrich MABE1126 | IF | 1:100 |
| Mouse anti phospho- γ H2AX | Merk, 05-636 | IF | 1:200 |
| Donkey anti rabbit IgG-A594 | Life Tech A21207 | IF | 1:500 |
| Goat anti mouse IgG-A647 | Life Tech ab136649 | IF | 1:500 |
| Rabbit anti UBF sera | In-house | ChIP | 60 µl/IP |
| Rabbit anti RPA127 sera | In-house | ChIP | 30 µl/IP |
| PAF53 | Proteintech #16145-1-AP | ChIP | 9.5 µl/IP |
| Mouse anti human CD45+PE Cy7 | BD 557748 | FACS | 1:500 |
| Mouse anti BrdU (clone 44) | BD 347580 | FACS | 1:500 |
| Mouse anti Ki67 | Leica Biosystem<br>NCL-L-Ki67-MM1 | IHC | 1:400 |
| Mouse anti alpha Smooth Muscle Actin | Sigma-Aldrich<br>A2547 | IHC | 1:10000 |

**Table S4.**
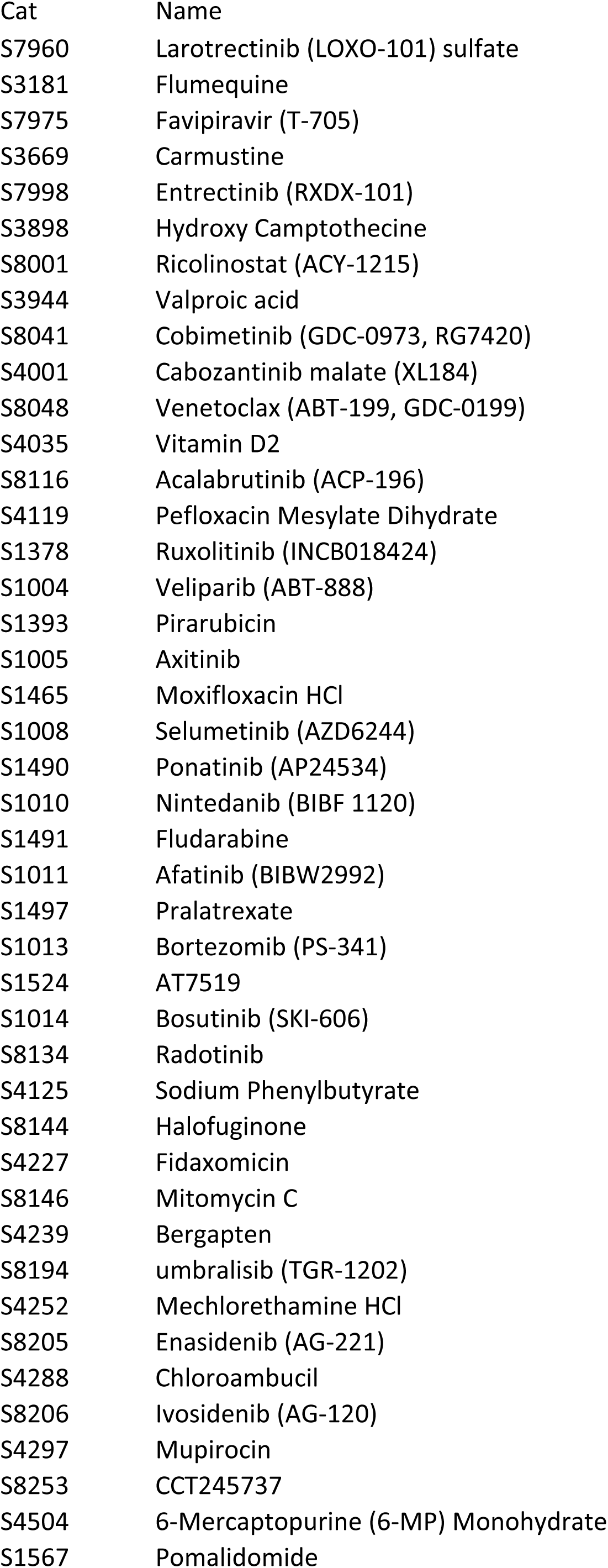
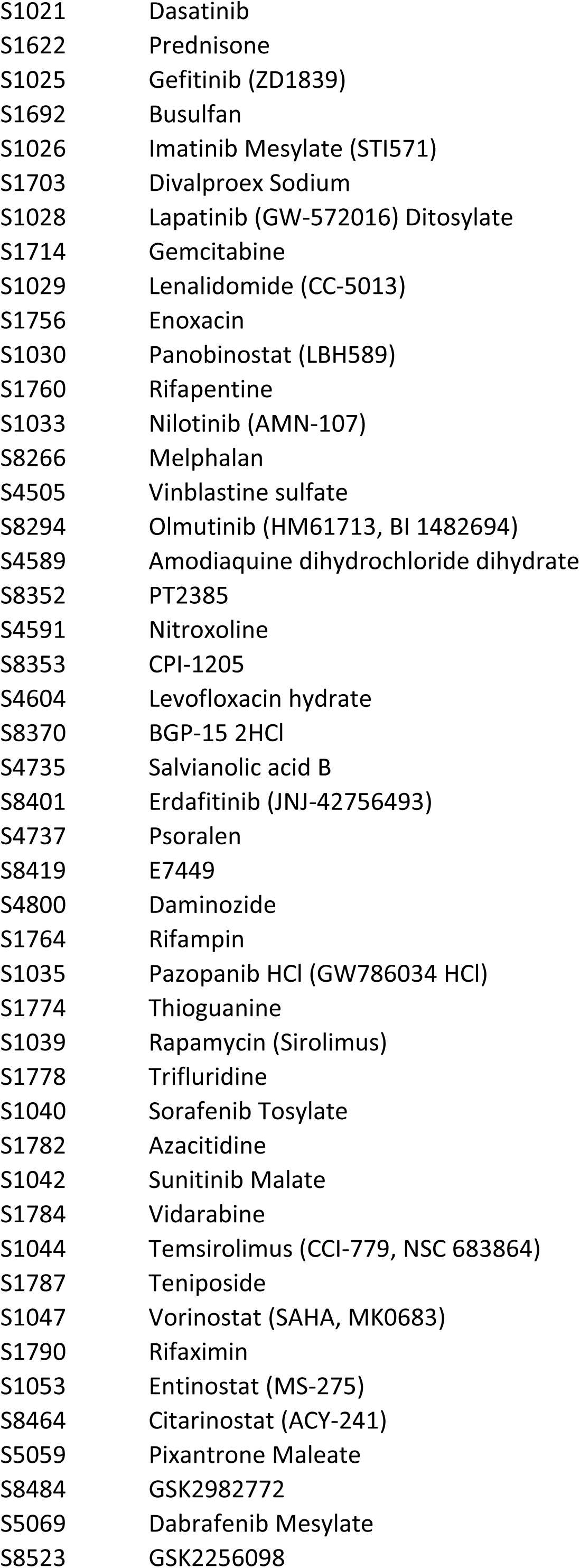
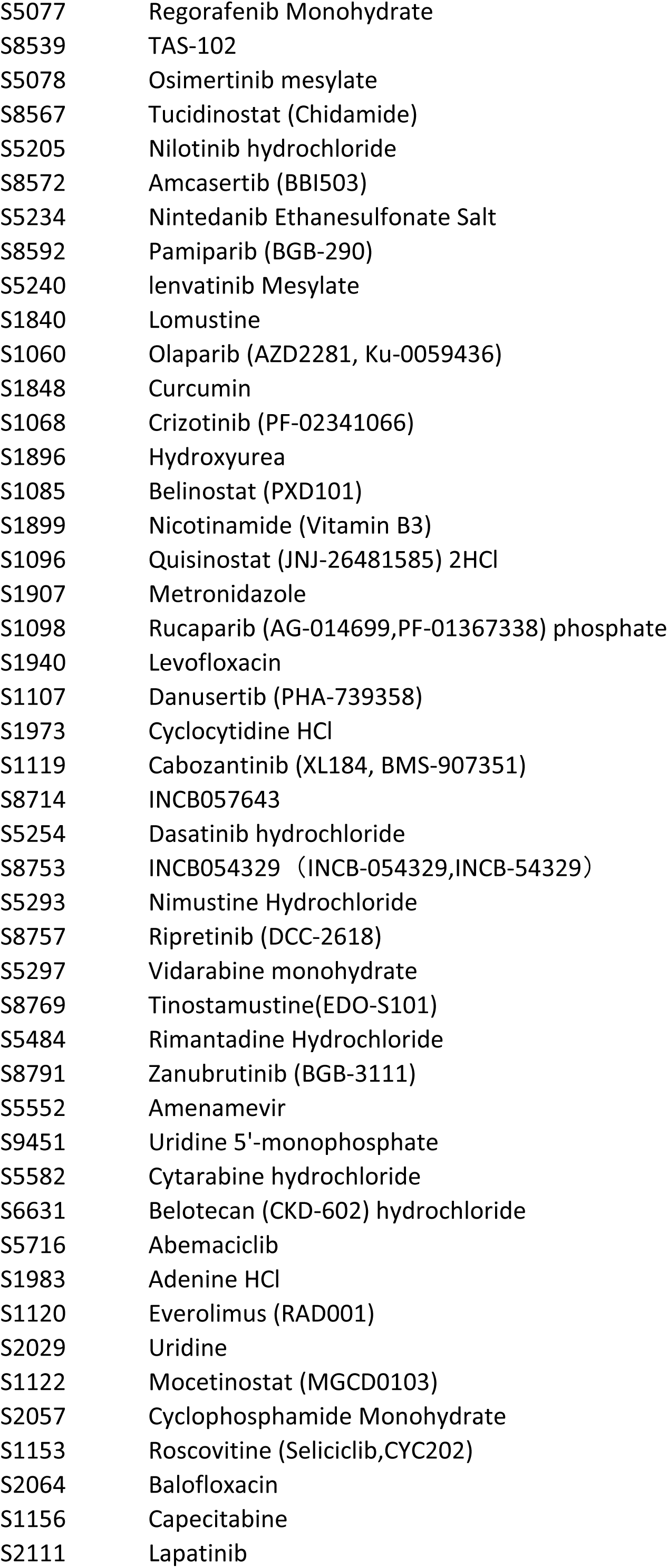
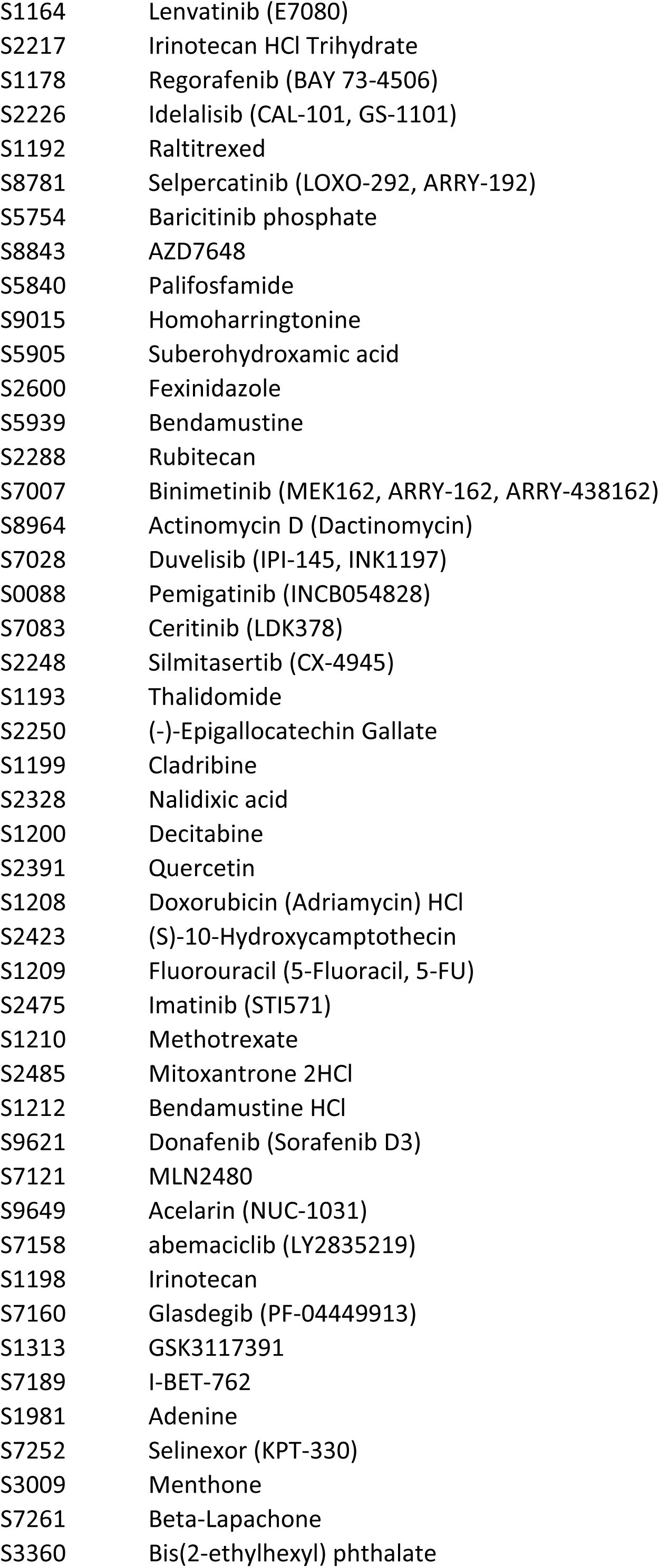
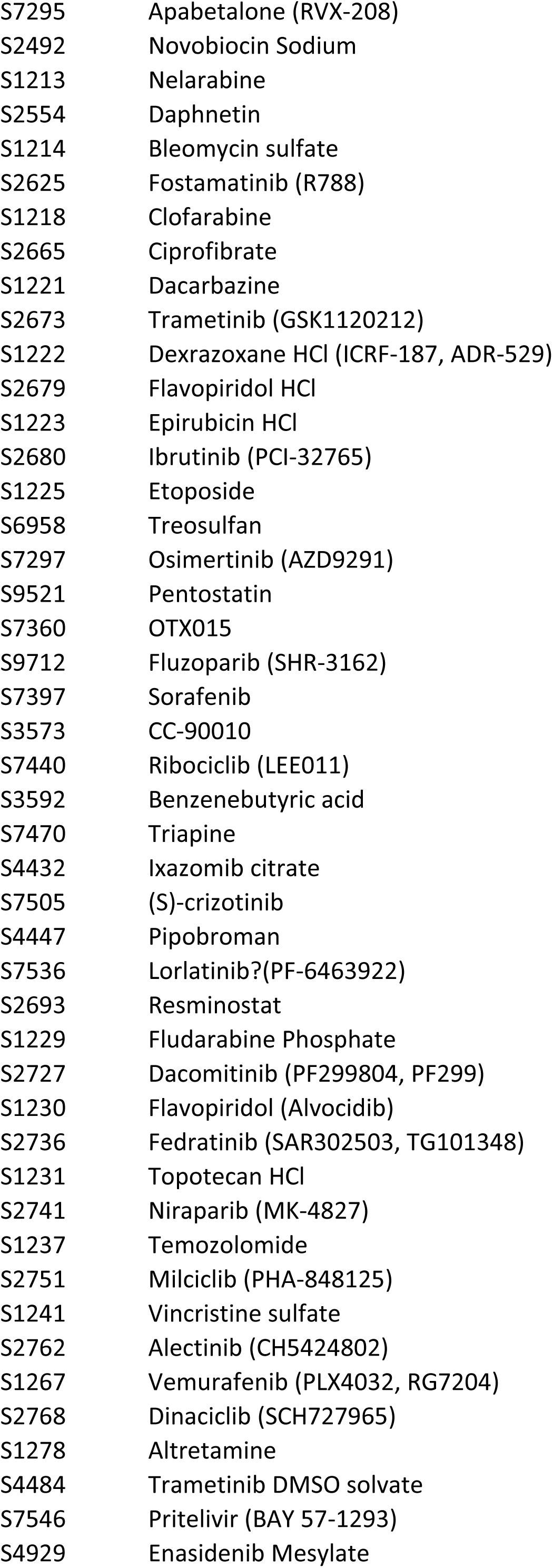
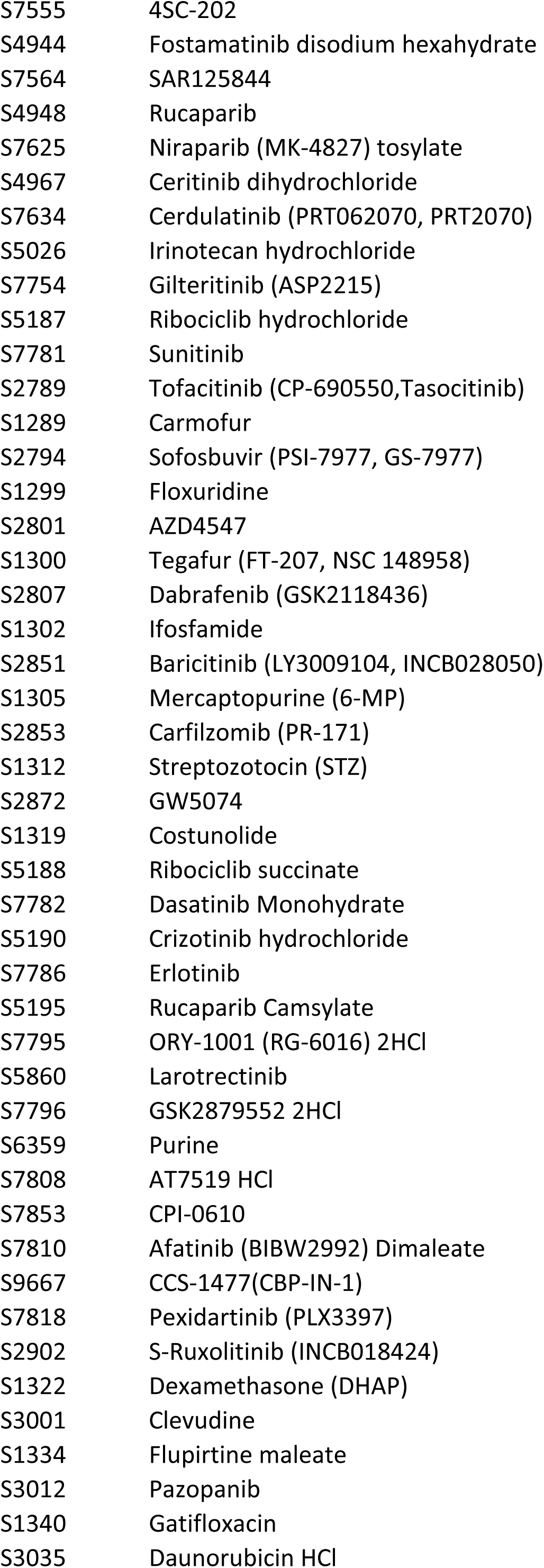
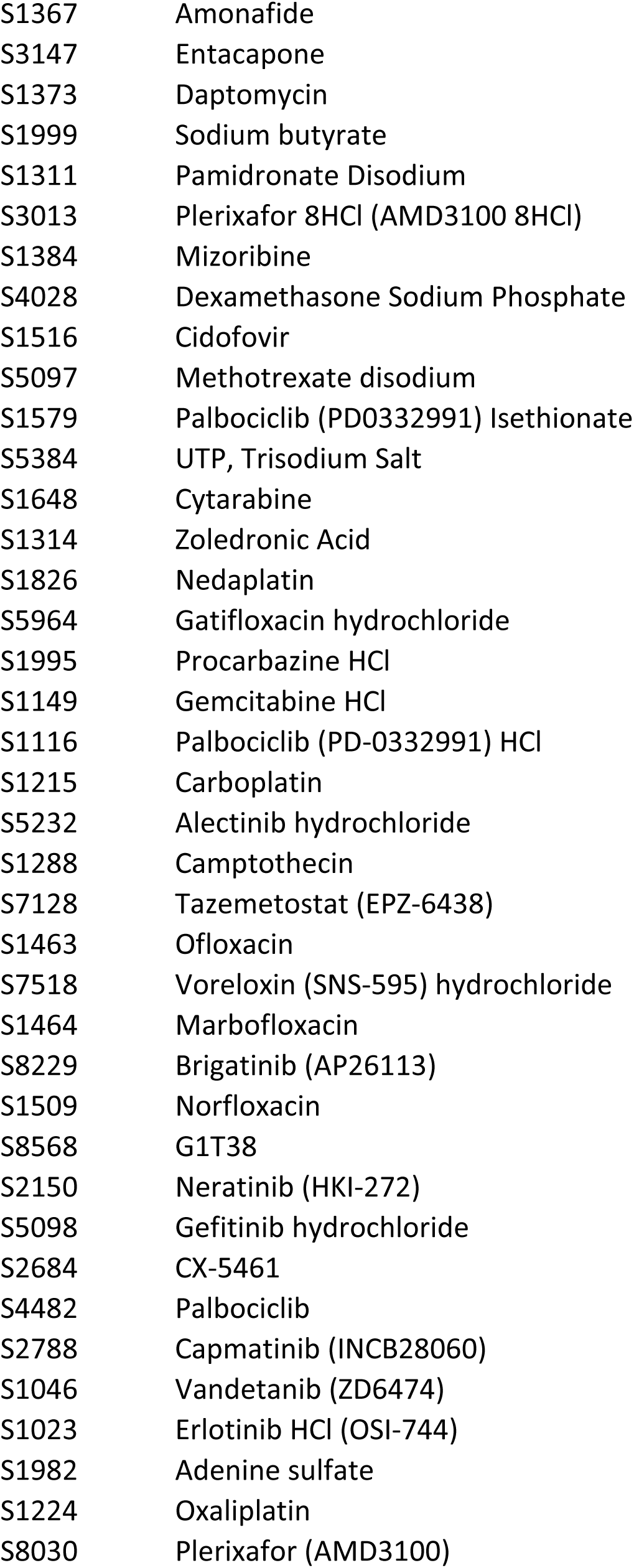

